# T cell transcription factor expression evolves as adaptive immunity matures in granulomas from *Mycobacterium tuberculosis*-infected cynomolgus macaques

**DOI:** 10.1101/2022.01.25.477732

**Authors:** Nicole L. Grant, Pauline Maiello, Edwin Klein, Philana Ling Lin, H. Jacob Borish, Jaime Tomko, L. James Frye, Alexander G. White, Denise E. Kirschner, Joshua T. Mattila, JoAnne L. Flynn

**Author notes:** Corresponding Author: JoAnne L. Flynn, Address: University of Pittsburgh, 5058 Biomedical Science Tower 3, 3501 Fifth Avenue, Pittsburgh PA, 15261. Competing Interests: Authors from the University of Pittsburgh and University of Michigan have no competing interests.

## Abstract

*Mycobacterium tuberculosis* (*Mtb)*, the causative agent of tuberculosis (TB), is a global health concern, yearly resulting in 10 million new cases of active TB. Immunologic investigation of lung granulomas is essential for understanding host control of bacterial replication. We identified and compared the pathological, cellular, and functional differences in granulomas at 4, 12, and 20 weeks post-infection in Chinese cynomolgus macaques. Original granulomas differed in transcription factor expression within adaptive lymphocytes with those at 12 weeks showing higher frequencies of CD8^+^T-bet^+^ T cells, while increases in CD4^+^T-bet^+^ T cells were observed at 20 weeks post-infection. The appearance of T-bet^+^ adaptive T cells at 12 and 20 weeks was coincident with a reduction in bacterial burden, suggesting their critical role in *Mtb* control. This study highlights the evolution of T cell responses within lung granulomas, suggesting that vaccines promoting the development and migration of T-bet^+^ T cells would enhance mycobacterial control.

## INTRODUCTION

*Mycobacterium tuberculosis (Mtb)*, the etiologic agent of tuberculosis (TB), has caused considerable morbidity and mortality for thousands of years (Barberis et al., 2017). Due to the COVID-19 pandemic, TB mortality is estimated to increase despite headway made in recent years by the End TB strategy (2020). There are still many unanswered questions about interactions of *Mtb* with its human host, and understanding these interactions are critical to development of improved treatments and preventive strategies. The complexity of TB disease is highlighted in the intricate pathological structure that forms following inhalation of *Mtb* bacilli, the lung granuloma. This dynamic structure is comprised of both innate and adaptive immune cells which undergoes cellular and molecular fluctuations throughout the course of infection leading to disparate trajectories, with some restricting or killing bacilli and others exhibiting a failure in bacterial control, propagating dissemination and progressive disease.

Despite extensive research, the immunological contributors to bacterial control in granulomas are not well understood. Studies investigating the role of adaptive T cells in *Mtb* infected mice have revealed IFN-γ dependent and independent mechanisms of control, suggesting that these cells have other critical functions in limiting TB disease (Kumar, 2017, Green et al., 2013, Gallegos et al., 2011). The influence of transcription factor (TF) expression, as a surrogate for immune cell function, has been studied in mice infected with *Mtb*, revealing a protective phenotype related to expression of the pro-inflammatory TF, T-bet (Sullivan et al., 2005). This is supported by studies in human patients with MSMD (Mendelian susceptibility to mycobacterial disease), which can be caused by defects in the genes encoding T-bet (*TBX21)* and RORγT *(RORC)* (Okada et al., 2015, Yang et al., 2020). A better understanding of TF expression within the context of the granuloma would enhance our ability to interpret T cell function in TB.

Investigating granuloma function in human TB disease is limited due to difficulty in obtaining representative samples, lack of data regarding time of infection, variable treatments, and little microbiologic information, necessitating the use of an animal model. Whereas many models for TB exist, non-human primates (NHPs) are invaluable as they reflect the range in granuloma pathology seen in humans (Capuano et al., 2003b, Lin et al., 2009). Cynomolgus macaques infected with virulent *Mtb* strains have been particularly useful as they recapitulate the full spectrum of infection outcomes seen in humans ranging from controlled (latent infection) to active TB disease (Lin et al., 2009, Lin et al., 2014, Maiello et al., 2017). We track granuloma formation following *Mtb* infection through positron emission tomography and computed tomography (PET CT) using ^18^F-fluorodeoxyglucose (FDG) as a PET probe, which incorporates into metabolically active host cells within granulomas (White et al., 2017, Coleman et al., 2014b, Coleman et al., 2014a, Lin et al., 2013). Serial PET CT scans over the course of infection provide a history for individual granulomas including time of detection, location, and changes in size and FDG avidity (Coleman et al., 2014b, White et al., 2017). Understanding the timing of granuloma formation is critical as granulomas observed at 4 weeks post-infection (termed original granulomas) represent those that are established by individual *Mtb* bacilli from the inoculum (Martin et al., 2017). Previous data indicate that granulomas which develop at later timepoints post infection (new granulomas), either through dissemination or slower growth of *Mtb*, have different features as they arise during an ongoing immune response, resulting in a decreased bacterial burden (Gideon et al., 2021, Lin et al., 2014). Differences in bacterial burden are also observed throughout the course of infection in granulomas; although the immune mechanisms responsible for this reduction in bacterial burden remain unclear, it likely relates to the evolving immunological state of the granuloma (Lin et al., 2014).

The present study is the first to investigate the interplay of TFs and bacterial dynamics in original NHP lung granulomas throughout the course of infection. We aimed to evaluate the cellular and functional changes in original granulomas over time using samples from macaques necropsied at early (4 weeks), mid (12 weeks), and late (20 weeks) timepoints post-infection. We observed temporal differences in TF expression in adaptive lymphocytes that correlate with bacterial burden, providing novel insights into the evolution of TB lung granulomas over time. These data suggest that protective adaptive immune responses are slow to develop and are coincident with a reduced bacterial burden in granulomas at later timepoints post-infection.

## RESULTS

### Study design and granuloma dynamics in original granulomas

To assess temporal changes in cellular composition, structure, and function of granulomas, 8 cynomolgus macaques were infected with a low dose of virulent *Mtb* and necropsied at 4 weeks (N=2), 12 weeks (N=3), or 20 weeks (N=3), representing early, mid, and late timepoints post-infection (Figure 1A). Granuloma formation was tracked over time using serial PET CT imaging starting at 4 weeks post-infection with PET CT scans performed every two to four weeks for the duration of the study. Based on these scans, we identified 94 original granulomas as those first observed on a 4-week scan which were individually harvested at necropsy and homogenized to a single cell suspension which was used for colony forming unit (CFU) quantification and flow cytometry. For granulomas >2mm, a portion was fixed in formalin for histology. In this study, our focus was on granuloma cellular composition, structure, and functional changes over time using only original granulomas. Consistent with previous data, original granulomas at 4 weeks post-infection had significantly higher CFU but similar size as measure by PET CT, when compared to original granulomas harvested at mid and late timepoints (Figure 1B) (White et al., 2017, Lin et al., 2014).

**Figure 1:**
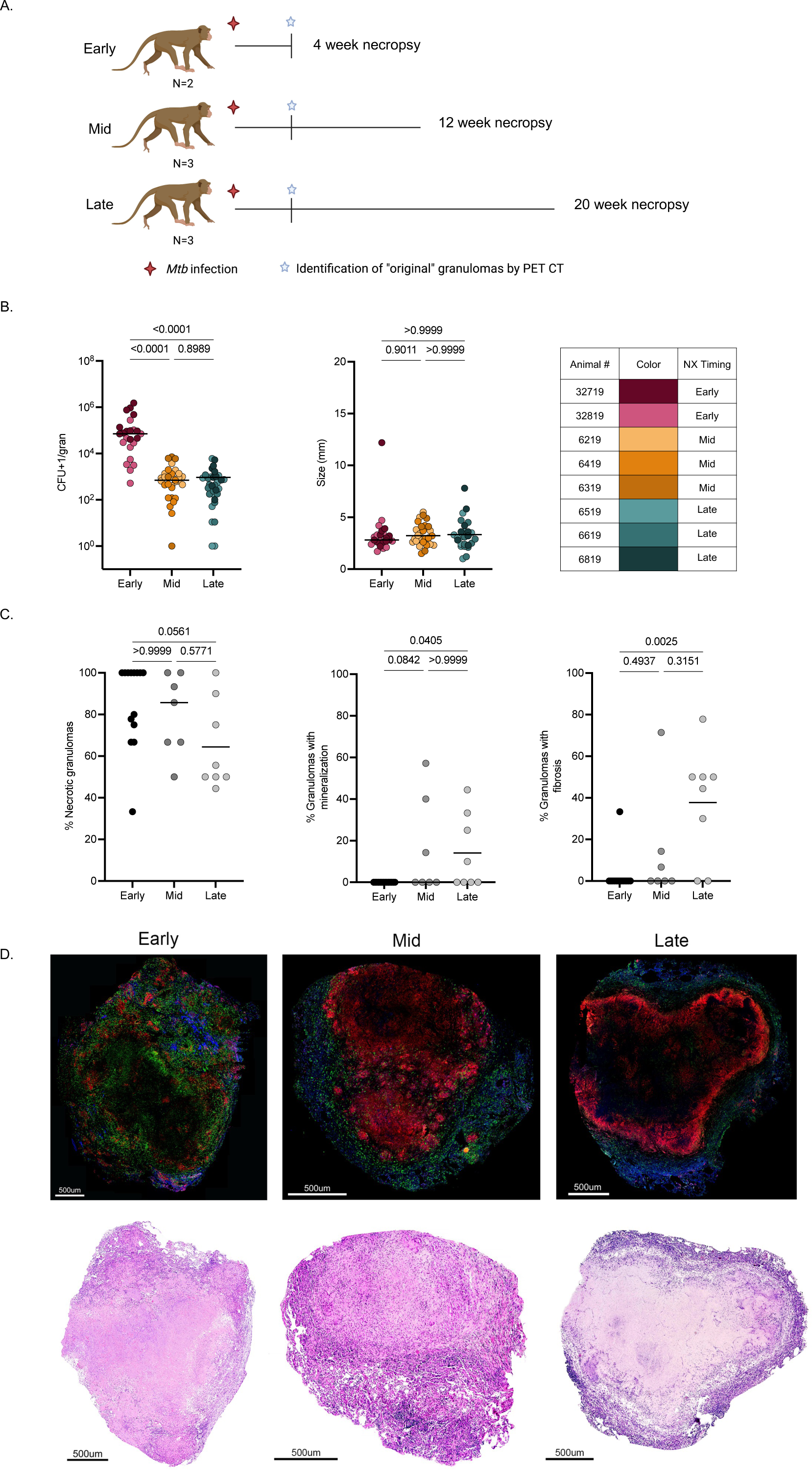
Study design, original granuloma dynamics and pathology over time. (A) Eight Chinese cynomolgus macaques were infected with low dose (10-19 CFU) *Mtb* Erdman and followed for 4 (early), 12 (mid), or 20 (late) weeks post-infection. Original granulomas were defined as those first observed on the 4 weeks post-infection PET-CT scan. (B) Individual granuloma bacterial burden [colony forming units (CFU)] and size (mm) from PET-CT. Individual points represent original lung granulomas (early n=24, mid n=31, late n=33). Table indicates each animal number and associated color in graphs. (C) Histologic characteristics of original lung granulomas in individual animals (frequency of each granuloma type) of banked samples at early, mid, and late timepoints categorized by H&E descriptions. Data points represent individual animals (N = 14 early, 7 mid, and 8 late animals). (D) Representative images of immunofluorescence staining for CD3 (green), CD11c (red), and CD163 (blue) with paired H&E staining of original granulomas from the early, mid, and late timepoints. For B and C statistics, Kruskal Wallis tests were performed with Dunn’s multiple comparisons-adjusted p values reported on the graphs.

### Histopathology and cellular spatial arrangement of original granulomas

Lung granulomas from *Mtb*-infected macaques are structurally similar to granulomas found in human TB patients (Flynn et al., 2015). Spatial arrangement is likely to be important for immune interactions and bacterial containment; thus, we compared the histological and spatial differences in original granulomas at early, mid, and late timepoints post-infection (Millar et al., 2021). In addition to samples from the monkeys dedicated to this study, we supplemented with banked samples of original granulomas isolated from monkeys at similar timepoints post-infection. The majority of original granulomas in individual animals had necrotic features, though lower frequencies of necrotic original granulomas were observed at the late timepoint compared to the early timepoint (Figure 1C). Necrotic or caseous features in granulomas can be observed in conjunction with other histologic findings such as fibrosis, collagenization, or mineralization (Flynn, 2011, Capuano et al., 2003b). While these secondary components were occasionally seen in original granulomas at the early timepoint, they were observed more frequently in original granulomas at the mid and late timepoints, suggesting they are temporally related to the transitional granuloma environment (Figure 1C). A range of histopathological classifications (as defined by our pathologist, EK) were also seen in original granulomas at all timepoints including classically necrotic, as well as rarer pathologies, such as evolving necrosis, early collagenization, fibrocalcific, and granuloma scarring (Supplemental Fig. 1C). We compared the CFU of fibrotic versus non-fibrotic granulomas from the mid and late timepoints and observed improved bacterial control in granulomas with fibrosis compared to non-fibrotic granulomas (Supplemental Fig. 1A), consistent with previous data (Lin et al., 2014).

While original granulomas at 12 and 20 weeks have reduced CFU compared to those at 4 weeks, sterilization of original granulomas from animals in this study and banked samples was rare at all timepoints, with the highest proportion at late timepoints (Supplemental Fig. 1B). Furthermore, sterile granulomas from the mid and late timepoints had split classifications based on histologic components, with some being fibrotic, neutrophilic, collagenic, or classically necrotic (Supplemental Fig. 1B). Classic necrotic granulomas are structurally organized, having a central caseous core surrounded by concentric rings of epithelioid macrophages and lymphocytes (Flynn et al., 2011, Flynn, 2011). With the exception of secondary structural elements (i.e. mineralization, fibrosis), visual comparison of H&E stained original necrotic granulomas from different timepoints revealed no distinct differences in overall granuloma structure (Figure 1D). We assessed this typical granuloma structure using immunofluorescent stained tissue sections for CD3^+^, CD11c^+^, and CD163^+^, finding more clusters of CD11c^+^ cells in granulomas from the early and mid timepoints. These clusters may be a precursor to the typically observed macrophage ring, suggesting that timing plays a role in the development of cellular spatial compartments in granulomas (Figure 1D).

### Cellular composition in original lung granulomas over time

Granulomas are dynamic structures composed of lymphoid and myeloid cells that contribute to bacterial control. We and others have previously reported substantial heterogeneity in granulomas in individual macaques and across macaques (Capuano et al., 2003a, Lin et al., 2014, Gideon et al., 2015, Maiello et al., 2017). Here we investigated whether there were differences in overall cellular composition in 94% of the original granulomas (88 of the 94) identified by PET CT using flow cytometry (Supplemental Fig. 2). A higher overall adjusted cell count (see Materials and Methods) was observed at the early timepoint for both myeloid and lymphocyte populations, whereas frequencies of these cell types were similar across all timepoints (Figure 2A and Supplemental Fig. 3A). There were no significant differences in the frequency of CD3^+^ cells across timepoints but significantly higher frequencies of CD20^+^ B cells at the mid timepoint and significantly higher frequencies of CD3^-^CD20^-^ cells (including NK and other innate lymphocytes) at the late timepoint (Figure 2C). Further analysis into CD3^+^ subsets revealed significantly lower frequencies of CD4^+^ T cells at the mid compared to the late timepoint (medians: mid-19.69%, late-24.39%, p value-0.0297) and significant differences in the frequencies of CD8^+^ T cells at each timepoint with the highest levels being at the mid timepoint (medians: early-17.36%, mid-27.46%, late-20.26%, p values-<0.0001 for early to mid, 0.0161 for mid to late, and 0.0112 for early to late) (Figure 2B-D). We observed low frequencies of both CD3^+^CD4^-^CD8^-^ and CD3^+^CD4^+^CD8^+^ cells at all timepoints with the highest median frequency for both cell types at the early timepoint (median: 4.81% and 4.95%, respectively) (Figure 2B and D).

**Figure 2:**
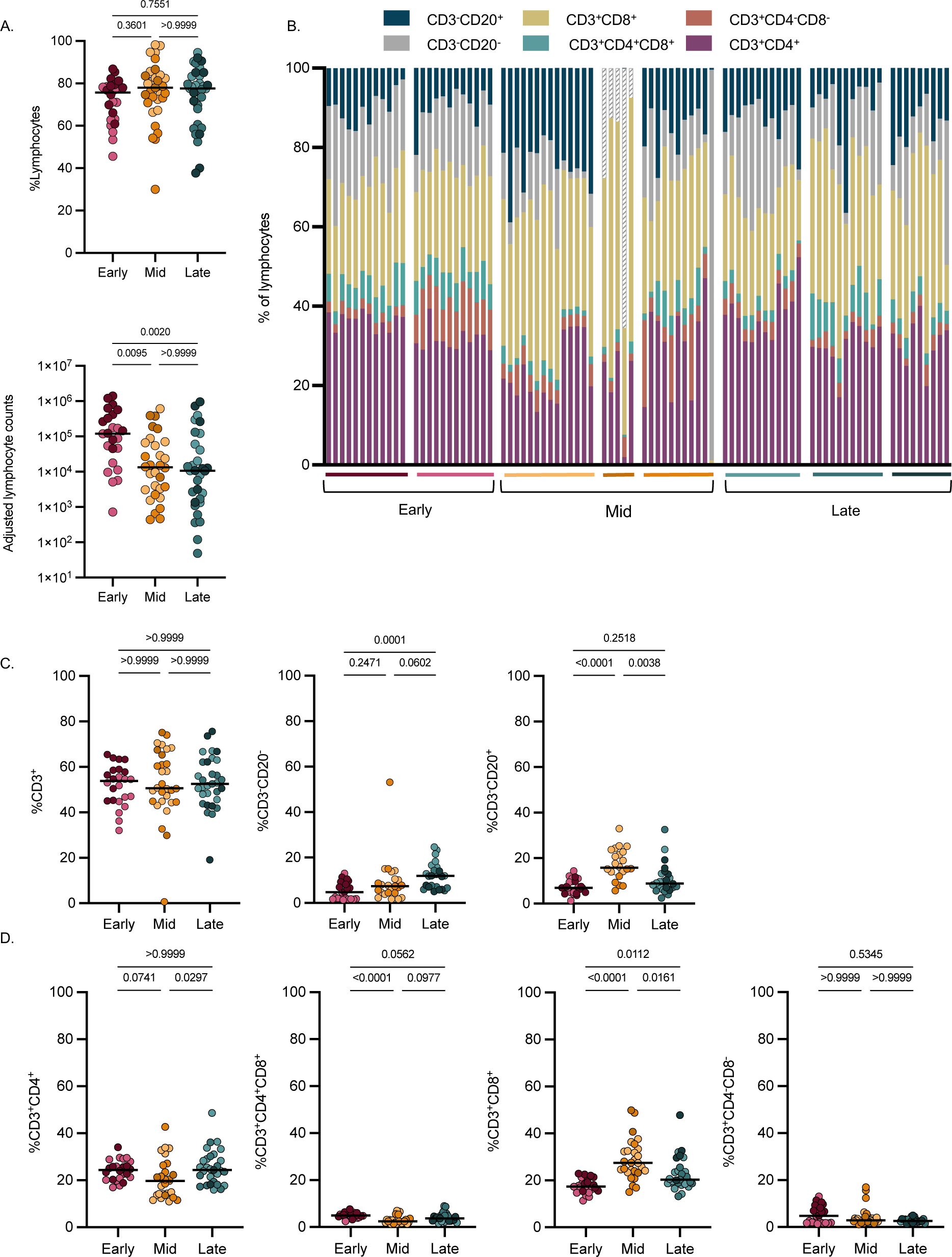
Few differences were observed in lymphocyte populations in original granulomas across timepoints. Original granulomas at each timepoint were evaluated for cellular composition by flow cytometry. (A) Frequency (top) and numbers (bottom) of lymphocytes. Each symbol is a granuloma. (B) Relative proportions of each cell type (% of lymphocytes) in individual granulomas. Each column is a granuloma and are separated by macaque (colored horizontal bar) and timepoint. (C) Frequency of CD3^+^, CD3^-^CD20^-^, and CD3-CD20^+^ of all live cells. (D) Frequency of CD4^+^, CD4^+^CD8^+^, CD8^+^ and CD4^-^CD8^-^ of all live cells. For A, C, and D statistics, Kruskal Wallis tests were performed with Dunn’s multiple comparisons-adjusted p values reported on graphs.

Myeloid cells make up approximately 25% of the cells in original granulomas regardless of timepoint (Supplemental Fig. 3A); since there were more cells in early granulomas these samples also had the highest adjusted myeloid cell count. At the early timepoint, there were higher frequencies of CD11c^+^, CD11b^+^, and CD163^+^ myeloid cells in granulomas (Supplemental Fig. 3B-C). To evaluate functional differences in myeloid cells we compared the frequency of these cells producing IL-10, TNF, and IFN-γ. While the frequencies of myeloid cells expressing any of the cytokines investigated was very low, there were significantly higher levels of IL-10 and IFN-γ in the late timepoint compared to the mid and early timepoints and significantly lower levels of TNF at the mid timepoint compared to early (Supplemental Fig. 3D).

### Transcription factor expression increases in granuloma adaptive T cells at 12 and 20 weeks post-infection

T cells are known to be important for controlling *Mtb*, however the breadth in function of these cells within lung granulomas remains incompletely studied. Our data (Figure 2C) and previous data suggest that although CD3^+^ cells make up approximately half of all cells in the granuloma (medians 50.63 - 53.78%) across timepoints, very few are reportedly producing cytokines within the granuloma (Gideon et al., 2015, Wong et al., 2018, Phuah et al., 2016, Millar et al., 2021). In this study, we analyzed lymphocytes directly from granulomas without additional stimulation to capture the functions that were occurring *in situ*, as granulomas contain *Mtb* antigens and live and dead bacilli which can serve as forms of T cell stimulation (Gideon et al., 2019). We observed very low frequencies of pro-inflammatory cytokine expression by all lymphocytes at any timepoint, though there is a significantly higher frequency of cells expressing IFN-γ and TNF in original granulomas from the early timepoint when compared to the mid and late timepoints (Supplemental Fig. 4A). Using markers that indicate activation, we observed the highest frequency of CD69^+^ lymphocytes at the early timepoint and the inverse for expression of PD-1 (Supplemental Fig. 4A) (Freeman et al., 2000, Ziegler et al., 1994, Cibrián and Sánchez-Madrid, 2017). This temporal expression of activation markers was similar in CD3^+^ subsets with PD-1 expression being highest at the late timepoint and CD69 at the early or mid timepoints (Supplemental Fig. 4B). The CD4^+^ and CD8^+^ adaptive T cells showed very low frequencies of IFN-γ and TNF, with slightly higher frequencies occurring at the early and mid timepoints for CD4^+^ T cells (medians: early-1.49%, mid-1.52%, late-0.25%) (Supplemental Fig. 4B). Notable was the high frequency of IFN-γ and TNF production from CD3^-^CD20^-^ cells at the early timepoint (medians: IFN-γ-16.85%, TNF-8.72%), emphasizing their early innate function (Supplemental Fig. 4C).

To provide additional insight into the function of granuloma cells we used flow cytometry to identify populations of transcription factor (TF) positive lymphocytes. The TF assessed were GATA3, T-bet, Foxp3, RORα and RORγT, which are regulators of cell differentiation and lineage commitment during an immune response for lymphoid cells (Nejati Moharrami et al., 2018, Saini et al., 2018, Yates et al., 2004, Wang et al., 2012, Fang and Zhu, 2017). Boolean gating indicated that the majority of lymphocytes in original granulomas at all timepoints did not express any of the TFs investigated (Figure 3A). Nevertheless, there was a significant increase in single and double TF expression at the mid and late timepoints post-infection (Figure 3A). When assessing only single TF expression in lymphocytes in granulomas across timepoints, there were statistically significant differences in all TFs investigated (Figure 3B-C). Although levels of Foxp3 and RORγT were very low at all timepoints, levels of RORα, GATA3 and T-bet were more substantial (Figure 3B-C).

**Figure 3:**
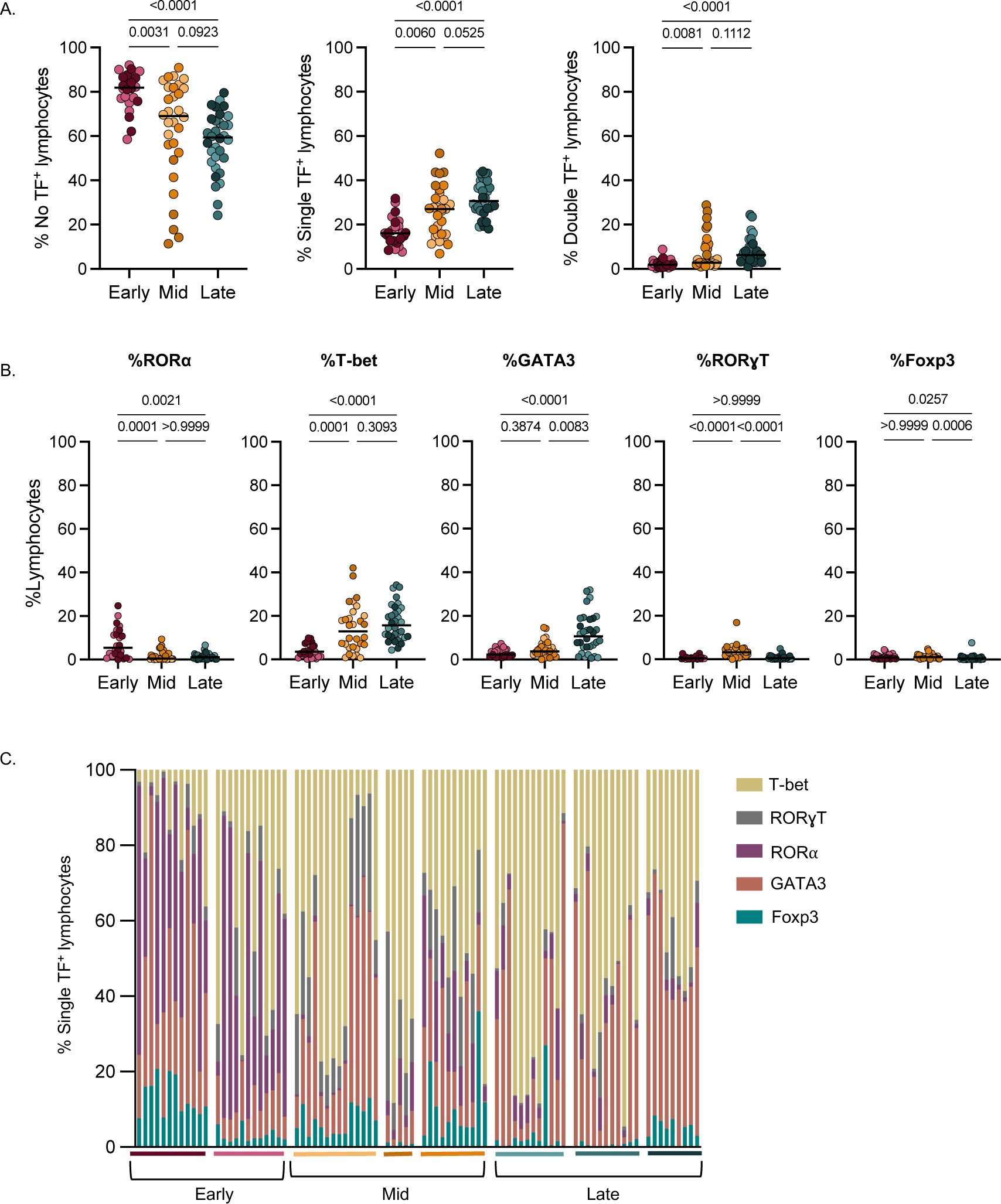
Temporal changes in transcription factor^+^ lymphocytes in original granulomas. (A) Original granulomas were assessed for transcription factor (TF) expression in lymphocytes via intracellular flow cytometry and Boolean gated to determine frequency of single TF expression and double TF expression in lymphocytes. (B) Frequency of lymphocytes expressing single TF from each timepoint. Individual symbols represent granulomas which are colored according to animal (Figure 1B). (C) Of the lymphocytes expressing a single transcription factor, the relative proportion of each of the five TF within individual granulomas is shown. Bars represent each granuloma and animals are noted by the colored horizontal line and grouped by necropsy timepoint. For A and B statistics, Kruskal Wallis tests were performed with Dunn’s multiple comparisons-adjusted p values reported on graphs.

Further analysis revealed that TF expression was dependent on lymphocyte subset. Conventional CD4^+^ T cells had a significant increase in the proportion of single TF^+^ cells in granulomas from the late timepoint (Supplemental Fig. 5A). Similarly, approximately 20% of CD4^+^CD8^+^ cells, which represent a subset of CD4 cells with a heightened activation profile, expressed a single TF at the early and mid timepoints, but had significantly higher expression at the late timepoint (Supplemental Fig. 5B) (Diedrich et al., 2019, Clénet et al., 2017). This contrasts with CD8^+^ T cells which had a median of 12.95% single TF expression at the early timepoint, with significantly higher frequencies of single TF^+^ cells at both the mid (median: 24.2%) and late (median: 25.3%) timepoints, revealing that increases in TF expression in CD8^+^ T cells occur earlier in granulomas compared to CD4^+^ cells (Supplemental Fig. 5A-C). CD3^+^CD4^-^CD8^-^ cells, which include NK T cells and some γδ T cells, had very low frequencies of single TF expression at the early timepoint and significantly higher frequencies at the mid and late timepoints with a highest median expression of 29.9% at the mid timepoint, although this expression was animal dependent (Supplemental Fig. 5D). There was a trend for higher single TF expression in B cells (CD3^-^CD20^+^) at the mid and late timepoints as compared to the early timepoint, although driven by one animal at the late timepoint (Supplemental Fig. 5E). Innate lymphocytes (CD3^-^CD20^-^ cells), however, had similar frequencies of cells with single positive TF expression at all timepoints (medians: 24.64%, 33.07%, and 25.35%) (Supplemental Fig. 5F). Conventional T cells (CD4^+^, CD4^+^CD8^+^, and CD8^+^) and B cells had significantly higher expression of two TFs at the late timepoint compared to the early and mid (Supplemental Fig. 5A-C, E).

To investigate specific TF expression within lymphocyte subsets, we compared single TF expression, keeping in mind the frequency of lymphocyte subsets in granulomas at each timepoint (number of granulomas assessed: early=24, mid=31, late=33) (Figure 4A). Consistent with what was observed in all lymphocytes (Figure 3B), there were relatively low frequencies of individual TF expression in many of the lymphocyte subsets at the early timepoint, with highest frequencies of RORα CD4^+^ T cells (Figure 3B-C, 4B). Innate lymphocytes (CD3^-^CD20^-^), had moderate levels of T-bet expression (medians: 16.7%, 27.92%, and 21.61%) at all timepoints investigated (Figure 4B), suggesting these cells play a role throughout infection. There was minimal TF expression in CD20^+^ B cells at all timepoints investigated, with modest increases in the frequency of RORα and GATA3 at the late timepoint, though these changes appear to be animal dependent (Figure 4B). At the early timepoint, we observed a low frequency of T-bet within adaptive CD3^+^ T cells (medians: CD4^+^:2.44%, CD8^+^:4.65%, and CD4^+^CD8^+^:7.5%) (Figure 4B). However, at 12 and 20 weeks post-infection there was a 5-fold increase in T-bet expression in CD8^+^ T cells (medians 22.2% and 21.2%). In contrast, there was no significant increase in T-bet^+^CD4^+^ T cells (including CD4^+^ T cells expressing CD8, i.e.CD4^+^CD8^+^) until 20 weeks post-infection (median CD4^+^:17.59% CD4^+^CD8^+^:21.0%) (Figure 4B) (Diedrich et al., 2019, Clénet et al., 2017). Although GATA3^+^CD4^+^ cells were rare in most granulomas, the proportions observed at the mid and late timepoints were modestly increased in some animals. While frequencies of cells expressing two TFs were low at all timepoints (<6% of lymphocytes, Figure 3A), we observed significant increases in the co-expression of T-bet^+^RORα^+^ and T-bet^+^GATA3^+^ at the late timepoint compared to the early and mid timepoints (Supplemental Fig. 5G). Taken together, these data support an evolution of the adaptive T cell response in granulomas over time.

**Figure 4:**
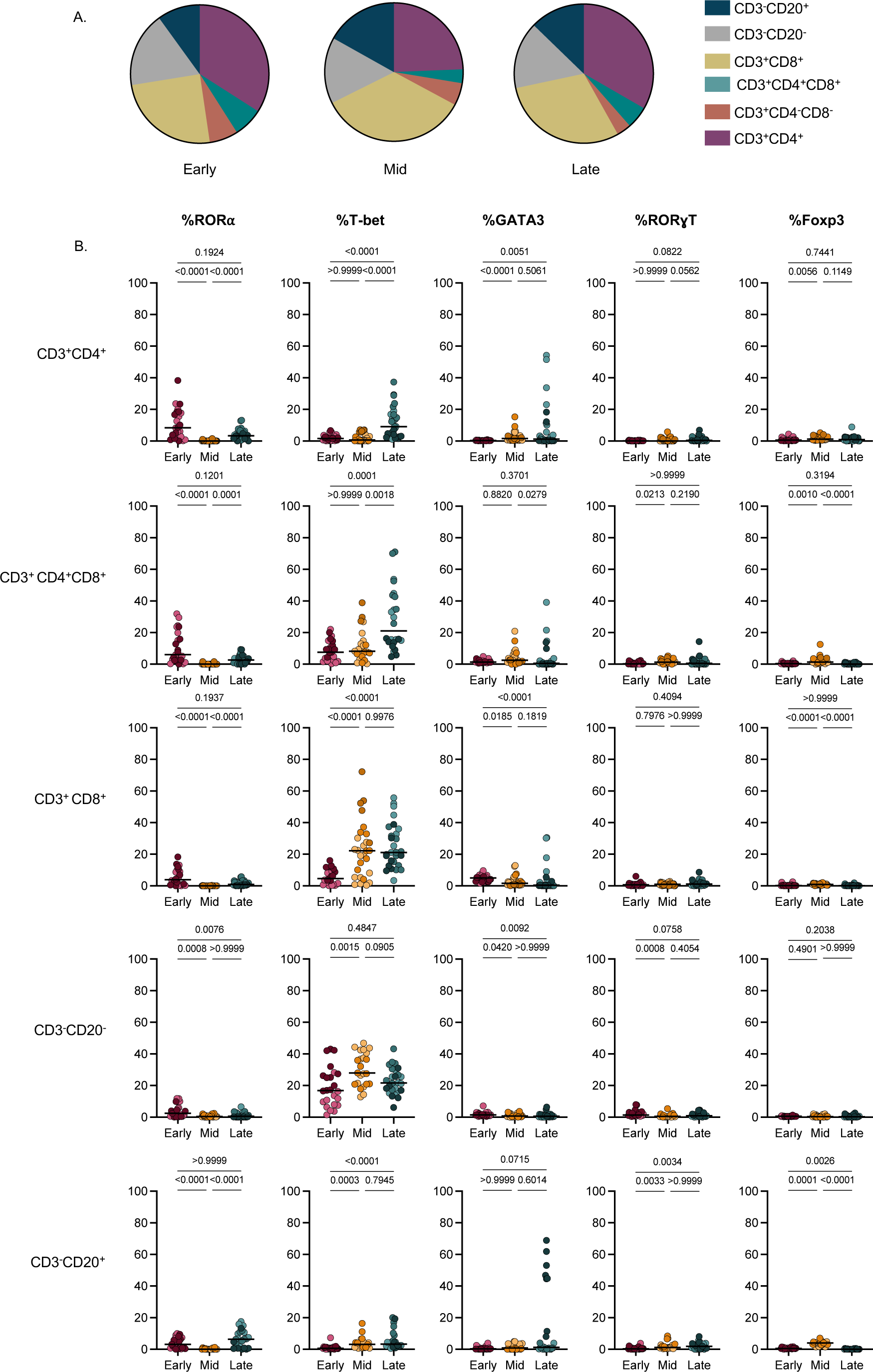
Delay in T-bet expression by conventional T cells in original granulomas. (A) Pie charts representing the average frequencies of denoted cell types (colored legend at right) from each timepoint (derived from Figure 2B). (B) Frequencies of each single transcription factor in lymphocyte subpopulations based on Boolean gated flow cytometry data. Kruskal Wallis tests were performed with Dunn’s multiple comparisons-adjusted p values reported on graphs.

To validate the flow cytometry data for TF expression and to investigate the localization of these cells in granulomas, we used immunofluorescence staining of granuloma tissue sections from the early, mid, or late timepoints post-infection. In keeping with the low levels of TF expression in early granulomas (Figure 4B), we found few CD3^+^ cells expressing RORα or GATA3 in granuloma tissue sections (Supplemental Fig. 6A, B). Although the frequency of Foxp3^+^ cells was consistently low at all timepoints, we could detect Foxp3 expressing CD3^+^CD4^+^ T cells (designated by arrows) (Supplemental Fig. 6C). Using a mid timepoint granuloma, we identified CD3^+^T-bet^+^ cells throughout the lymphocyte region as well as within clusters of CD11c^+^ macrophages (macrophage region), suggesting a potential interaction between these cell types (Figure 5A). There was consistent nuclear localization of T-bet (designated by arrows), indicating an activated cellular phenotype (Figure 5A) (McLane et al., 2013).

**Figure 5:**
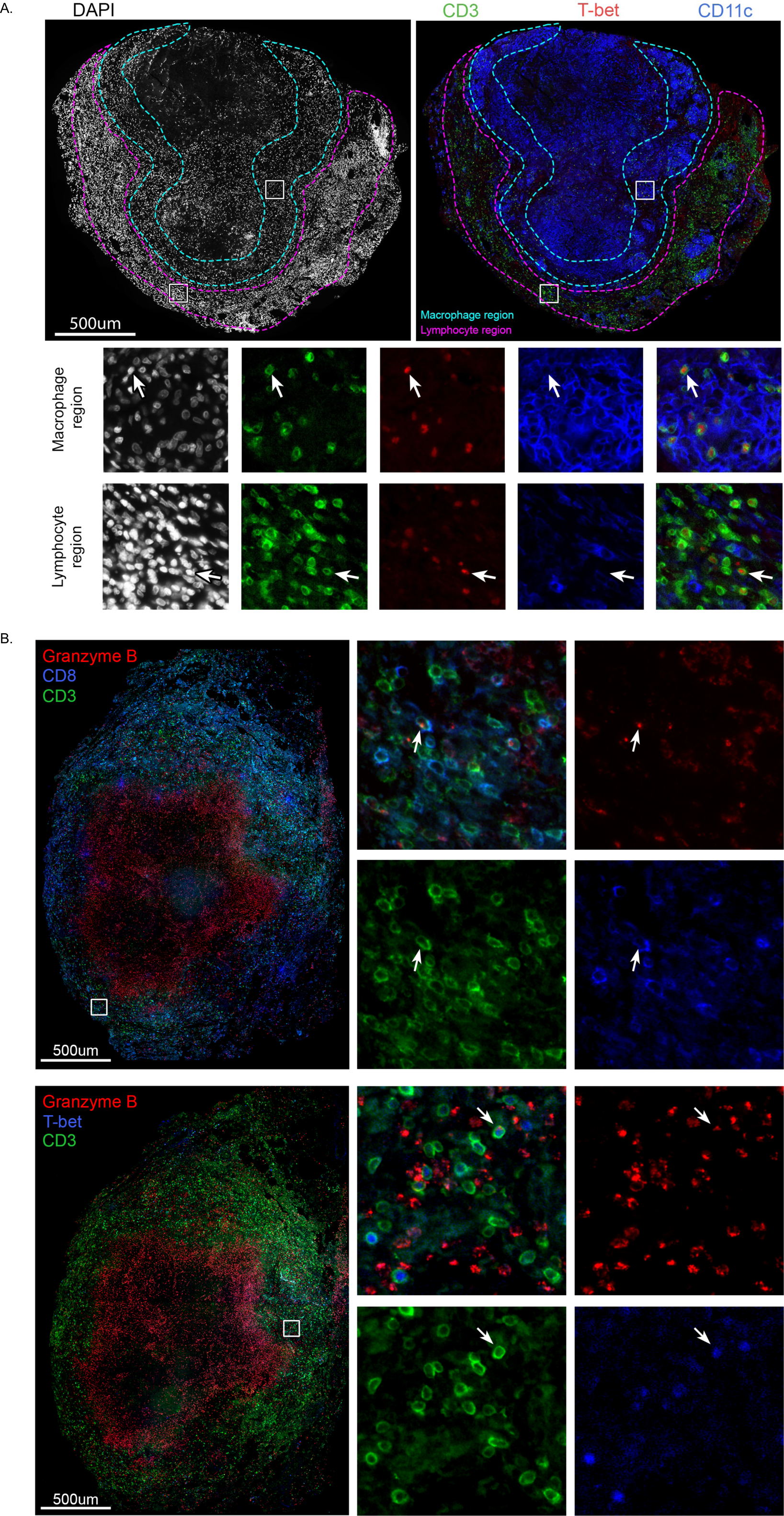
Presence, localization, and function of T-bet^+^ lymphocytes in original granulomas. (A) Original granuloma from an animal 12 weeks post-infection with Dapi staining (left panel), CD3 (green), T-bet (red), and CD11c (blue) staining (right panel). Insets as denoted in large image show localization of T-bet^+^CD3^+^ cells within macrophage regions (teal) as well as the lymphocyte cuff (magenta). (B) Original granuloma isolated from an animal 12 weeks post-infection showing localization of granzyme B (red) in CD3^+^ (green) CD8^+^(blue) cells (co-registered as teal and denoted with arrows) (top panel) and CD3^+^(green)T-bet^+^(blue) expressing granzyme B (red) (denoted by arrows) (lower panel).

### Transcription factor positive cells have higher frequencies of pro-inflammatory cytokines than transcription factor negative cells

To assess the functionality of the TF^+^ cells we compared the frequency of pro-inflammatory cytokine expression in TF^+^ cells versus TF^-^ cells from the same sample (Figure 6A-C). At all timepoints, RORα^+^ or T-bet^+^ CD4^+^ and CD4^+^CD8^+^ T cells had higher frequencies of IFN-γ, TNF and CD69 expression compared to TF^-^ cells, supporting that TF^+^ cells are activated and functional (Figure 6A, B). T-bet^+^CD4^+^ and T-bet^+^CD4^+^CD8^+^ T cells had a higher frequency of PD-1 at the mid and late timepoints, suggesting that these cells have a different activation profile than RORα^+^CD4^+^ cells at the early timepoint. T-bet^+^ innate lymphoid cells (CD3^-^CD20^-^) at the early timepoint had significantly higher production of both IFN-γ and TNF compared to T-bet^-^ cells, indicating that T-bet^+^ innate lymphocytes are one contributor of pro-inflammatory cytokines in early granulomas (Supplemental Fig. 7).

**Figure 6:**
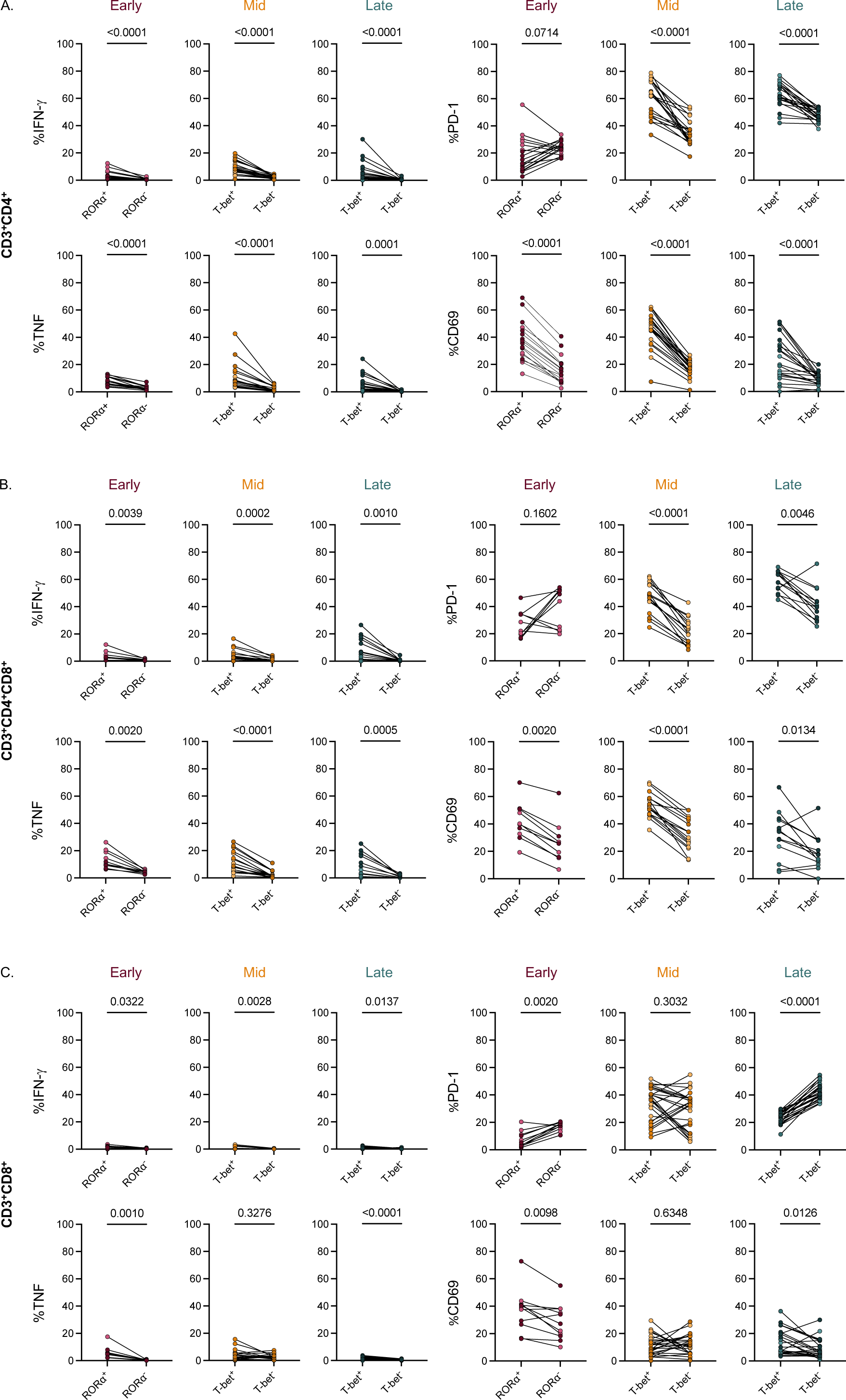
Transcription factor^+^ cells produce more cytokines than transcription factor^-^ cells in original granulomas at all timepoints. Comparison of pro-inflammatory cytokine (IFN-γ, TNF) production or activation marker (CD69, PD-1) expression in TF^+^ T cells (RORα for early, T-bet for late) compared to TF^-^ T cells for CD4+ T cells (A), CD4^+^CD8^+^ T cells (B), and CD8^+^ T cells (C). All comparisons were performed using Wilcoxon signed-rank tests.

CD8^+^ T cells showed very low frequencies of cytokine^+^ cells at all timepoints. Despite this, TF^+^CD8^+^ cells exhibited higher frequencies of pro-inflammatory cytokines when compared to TF^-^ CD8^+^ cells (Figure 6C). TF expression in conjunction with low frequencies of cytokine expression suggested that CD8^+^ T cells are contributing to the granuloma environment through other functions, such as producing cytotoxic molecules. To investigate this, we stained a granuloma from 12 weeks post-infection, observing granzyme B localization within CD3^+^CD8^+^ cells (designated by arrows) (Figure 5B). We observed colocalization of granzyme B with CD3^+^T-bet^+^ cells in the same granuloma and in a late timepoint granuloma (arrows) (Figure 5B, Supplemental Fig. 6D). The available CD8 and T-bet antibodies could not be used in tandem for staining, limiting the ability to directly identify granzyme B expression in CD8^+^T-bet^+^ cells but, taken together, our data suggests that CD3^+^CD8^+^ cells expressing T-bet produce granzyme B at 12 weeks post-infection. Of note, we observed granzyme B staining within the caseum; although this was not associated with intact nuclei (Dapi), this may be true signal, potentially suggesting the caseum is a sink for granzyme B. The presence of granzyme B within the caseum was also observed in granulomas from the late timepoint but varied in abundance between granulomas (Figure 5B, Supplemental Fig. 6D).

### Frequency of T-bet^+^ cells negatively correlates with granuloma bacterial burden

Granulomas can contribute to *Mtb* protection by promoting immune responses that kill or restrict bacterial replication; conversely, they may promote disease by supporting *Mtb* growth and dissemination. By analyzing snapshots of granulomas at different timepoints we can begin unravelling specific immune elements that are associated with a reduction in bacterial burden. One striking difference between granulomas at 4 weeks versus those at 12 and 20 weeks is the presence of T-bet^+^ T cells. Correlation analyses revealed a modest but significant negative correlation between the proportion of all T-bet^+^ lymphocytes, CD4^+^T-bet^+^, or CD8^+^T-bet^+^ cells and CFU per granuloma (Figure 7A-C). This association suggests that T-bet^+^ lymphocytes are one contributor to the reduction in bacterial burden seen in granulomas at 12 and 20 weeks post-infection.

**Figure 7:**
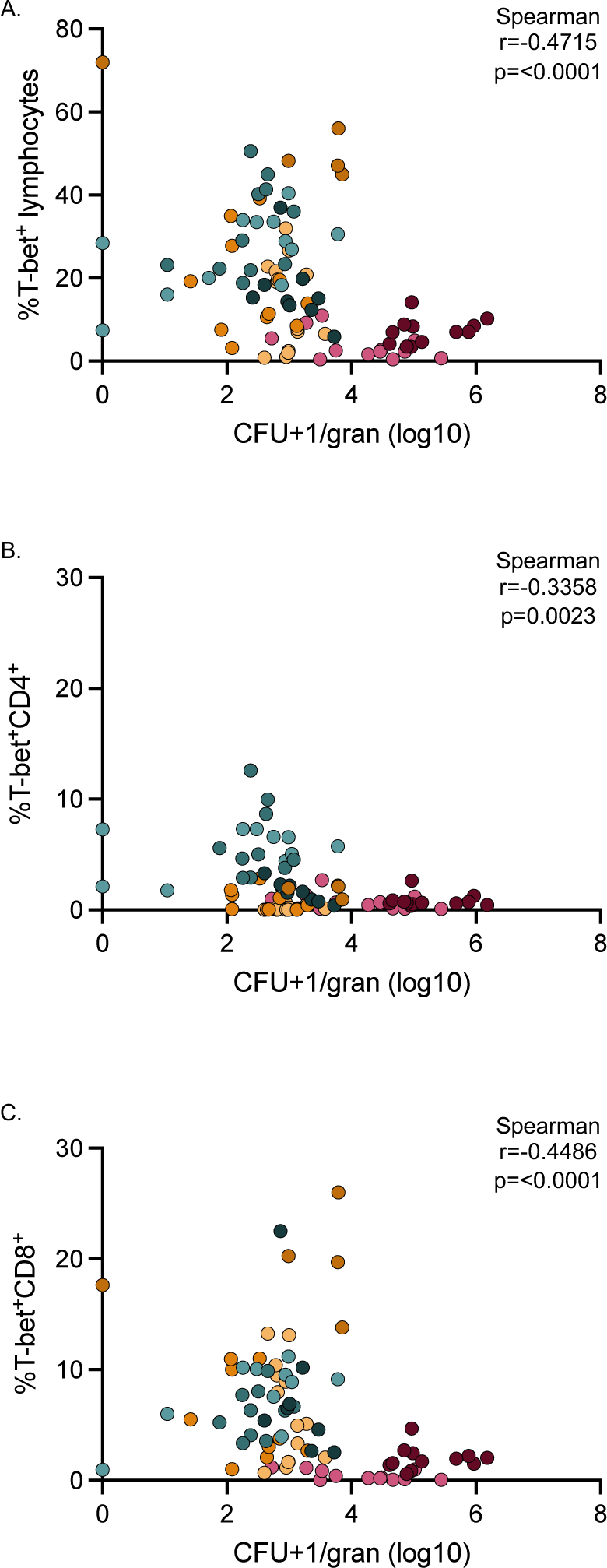
Bacterial burden negatively correlates with frequencies of T-bet^+^ lymphocytes. (A) Frequency of T-bet^+^ lymphocytes vs. log10 CFU per granuloma. (B) Frequency of T-bet^+^CD4^+^ vs. log10 CFU per granuloma. (C) Frequency of T-bet^+^CD8^+^ T cells vs. log10 CFU per granuloma. CFU per granuloma was log transformed after adding 1 (to avoid having undefined values). Non-parametric Spearman correlation analyses were performed with rho and p values noted on individual graphs.

## DISCUSSION

Understanding the process of bacterial restriction and containment in lung TB granulomas is critical for identifying new targets for vaccines and therapeutics. Here we compared original granulomas, i.e. those that arise from initial infection as determined by PET CT, from three distinct timepoints (4, 12, and 20 weeks) post-infection in NHPs (Martin et al., 2017). These timepoints represent early infection (early), the beginning of infection control (mid), and late infection (late), respectively. This affords an opportunity for temporal analysis of granuloma structure, cellular composition, and function, providing insight into some of the immune components that contribute to a reduction in bacterial burden. Our results reveal that immune responses mediated by T-bet expressing T cells are delayed in granulomas, supporting that adaptive immunity in granulomas evolves over time. The slow evolution of adaptive immunity likely contributes to the ease of establishment of *Mtb* infection and substantial growth of the pathogen in early granulomas, where one originating bacillus in a granuloma reaches ∼10^5^ CFU by 4 weeks (Lin et al., 2014). This suggests that vaccine-mediated enhancement of CD8^+^ T cell responses, in addition to CD4^+^ T cell responses, and rapid recruitment to the airways and lung following infection could improve protection against infection and progressive disease. Recent reports of vaccine strategies using CMV producing *Mtb* antigens, or intravenous and mucosal BCG support this concept as they induce strong CD4^+^ and/or CD8^+^ T cell responses in airways and lungs and induce robust protection against *Mtb* infection or disease in macaques (Darrah et al., 2020, Hansen et al., 2018, Dijkman et al., 2019).

Granuloma structure has been investigated in various animal models and in humans, revealing a wide range of histopathological features. In humans and NHPs infected with *Mtb*, granulomas are classified as having necrotic (caseous), fibrotic, non-necrotic, mineralized, scarring, cavitary, and suppurative phenotypes (Flynn et al., 2015, Flynn, 2011, Lin et al., 2014). Necrotic granulomas were the most observed phenotype among original granulomas in this study, regardless of timepoint post-infection, however, a range of granuloma structures were observed in original lesions at the mid and late timepoints including the presence of fibrosis. Prior studies in NHPs have investigated the role fibrosis plays in granuloma healing and containment which is consistent with the lower CFU in original granulomas with fibrosis observed in this study (Warsinske et al., 2017).

There were no major differences in the proportion of lymphocytes or myeloid cells in granulomas at the different stages of infection. There was a higher frequency in myeloid cell expression of the integrins CD11c and CD11b that aid in adherence, migration, and phagocytosis at the early timepoint (Lukácsi et al., 2020). A similar trend was observed in the frequency of cells expressing the scavenger receptor CD163, a marker often observed on alveolar macrophages (Bharat et al., 2016). While there are several potential hypotheses for these differences, one possibility is that granulomas from distinct timepoints are comprised of different myeloid subsets: i.e. early granulomas have more neutrophils and alveolar macrophages and fewer epithelioid macrophages. This hypothesis is reasonable as studies investigating the early events in granuloma formation and pathogen clearance reveal involvement of alveolar macrophages and neutrophils in the phagocytosis of bacteria and in bacterial clearance, respectively (Cohen et al., 2018). Regarding myeloid cell functionality, in late granulomas, we observed modest frequencies of cells producing IFN-γ or IL-10. As levels of IL-10 are higher than IFN-γ, myeloid cells may be more anti-inflammatory at late timepoints, possibly modulating pathology or contributing to *Mtb* persistence (Redford et al., 2011).

It is generally accepted that T cells are critical for the control of *Mtb* through production of cytokines such as IFN-γ, although it is not the only mediator of protection and additional T cell functions are likely to be of equal importance (de Martino et al., 2019, Gideon et al., 2015, Lin et al., 2012, Gallegos et al., 2011). In fact, vaccines that induce production of IFN-γ by CD4^+^ T cells are not always successful in the prevention of TB in animal models or humans (Darrah et al., 2019, Darrah et al., 2020, Verreck et al., 2009, Tameris et al., 2013, Abou-Zeid et al., 1997, Orr et al., 2015, Griffiths et al., 2016). Our objective was to investigate lymphocytic phenotypes in granulomas at different timepoints not only for cytokine production but also for broader functionality using TFs.

The role of TFs as lineage specifying amongst T cells has been well established, connecting the expression of T-bet, GATA3, Foxp3, RORα, and RORγT to T_H_1, T_H_2, T_reg_, and T_H_17 cells, respectively (Szabo et al., 2000, Ivanov et al., 2006, Zheng and Flavell, 1997, Hori et al., 2003). In granulomas, regardless of timepoint, the majority of lymphocytes were not expressing any of these TFs. One possible explanation for low levels of TF expression is that lymphocytes in granulomas rarely encounter antigen presenting cells infected with *Mtb*, due to the spatial localization of cells in the granuloma, which has been predicted in previous studies using mathematical modeling (Millar et al., 2021). Another hypothesis is that many of the lymphocytes in granulomas are not specific for *Mtb* but migrate to the granuloma due to inflammatory signals and chemokines from infected or activated cells (Gideon et al., 2015, Wong et al., 2018, Millar et al., 2021). The necessity and function of adaptive T cell recruitment to *Mtb* infected lungs has been investigated using various models, with studies in mice highlighting the roles of IL-23 and IL-17 as critical in the early recruitment of functional T cells to the lungs (Marino et al., 2011, Millar et al., 2021, Kauffman et al., 2018, Domingo-Gonzalez et al., 2017, Khader et al., 2007). While contributing events in the evolution of the adaptive immune response are not fully understood, bacterial reduction likely depends on spatially positioned, functional T cells for interaction with infected macrophages, as observed here with T-bet^+^ cells in granulomas at 12 and 20 weeks post-infection.

At the 4 week timepoint, we observed two pronounced phenotypes: 1) production of IFN-γ and TNF by innate lymphocytes and 2) RORα expression in lymphocytes. RORα is a member of the retinoid orphan receptor family and canonically known for its role in the development of ILCs and Th17 cells (Yang et al., 2008, Ferreira et al., 2021, Lo et al., 2019). More recently, RORα was shown to be expressed in activated CD4^+^ T cells of T_H_1 and T_H_2 helper cell lineages with relationships to chemotaxis and cell migration (Haim-Vilmovsky et al., 2021). Studies in *Mtb* infected humans and mice also report an accumulation of ILCs in infected lungs and a protective role of ILC3s expressing the RORα homolog, RORγT, at early timepoints post-infection (Ardain et al., 2019). This suggests that innate lymphocytes as well as lymphocytes expressing ROR family proteins are vital cells in the early phases following *Mtb* infection, possibly for recruiting additional cells into the lung or granuloma which facilitate bacterial killing at later timepoints.

At 12 weeks post-infection, there was a substantial increase in the frequency of T-bet expression in CD8^+^ T cells followed by the increased frequency of CD4^+^T-bet^+^ cells at 20 weeks post-infection. The presence and role of T-bet has been investigated in the context of several infectious diseases, including TB, revealing its critical function in controlling infection through production of pro-inflammatory mediators, T cell trafficking, and inhibition of other T cell fates (Pritchard et al., 2019, Szabo et al., 2000, Sullivan et al., 2005, Lazarevic et al., 2013, Lord et al., 2005). The earlier temporal increase in T-bet^+^CD8^+^ T cells which is coincident with a reduction in bacterial burden suggests these cells play a crucial role in bacterial control. This is consistent with our recent single cell RNAseq data from NHP granulomas at 10 weeks post-infection (Gideon et al., 2021). Despite increases in T-bet^+^CD8^+^ cells at the mid and late timepoints, low frequencies of these cells produce pro-inflammatory cytokines. We instead observed granzyme B expression in these cells, suggesting a more traditional cytotoxic function. At the late timepoint, there are higher frequencies of CD4^+^T-bet^+^ and CD4^+^CD8^+^T-bet^+^ cells producing IFN-γ or TNF and expressing PD-1 and CD69 when compared to T-bet^-^ cells in the same granuloma. Surface expression of PD-1 is associated with activated cells or functional deficiency when co-expressed with other exhaustion markers, particularly in the context of cancer and viral infection (Barber et al., 2006, Dong et al., 2019). Studies in mice have identified an *Mtb* specific subset of PD-1^+^CD4^+^ T cells that are functional and highly proliferative, potentially acting as a self-replenishing source of CD4^+^ T cells in TB (Reiley et al., 2010). We have consistently observed low levels of cytokine production from T cells in NHP granulomas, which initially suggested an exhausted phenotype. However, we previously reported low levels of exhaustion markers on CD3^+^ T cells and no difference in the cytokine production in cells with or without specific exhaustion markers (Wong et al., 2018). These studies, taken together, suggest that PD-1 expression in T-bet^+^CD4^+^ cells is likely related to T cell activation or regulation rather than exhaustion. Furthermore, our staining for T-bet^+^ cells in granulomas showed nuclear localization of T-bet, which is indicative of an activated rather than exhaustive phenotype (McLane et al., 2013, McLane et al., 2021).

Though this study offers insight into the evolving granuloma environment, our flow panels for this experiment did not include antibodies to detect cytotoxic effector molecules or additional transcription factors which limited our ability to comprehensively assess immune responses in granulomas. Future studies will include assessment of additional effector molecules with a focus on cytotoxic effectors such as granzymes and granulysin. In addition, applying single cell RNA sequencing on cells isolated from granulomas at distinct timepoints post infection will provide a more robust and unbiased approach, corroborating the data provided herein.

In this study, early granulomas were characterized as having higher CFUs accompanied by higher frequencies of innate lymphocytes producing inflammatory cytokines and lower frequencies of adaptive lymphocytes expressing T-bet, or any of the TFs investigated. The increase in T-bet^+^ CD8^+^ T cells preceded the appearance of T bet^+^ CD4^+^ T cells at the later timepoints. This suggests that 4 weeks post-infection is prior to the development of an adaptive immune response, whereas at 12 weeks (CD8^+^) and 20 weeks (CD4^+^ and CD8^+^) functional adaptive lymphocytes appear in granulomas, highlighting the prolonged time frame needed for development of a robust adaptive T cell response to *Mtb* (Mehra et al., 2010). When comparing original granulomas, the proportion of T-bet^+^ lymphocytes, T-bet^+^CD4^+^ and T-bet^+^CD8^+^ cells negatively correlated with bacterial burden, suggesting that conventional T cells (CD4^+^ and CD8^+^) expressing T-bet contribute to the reduction in bacterial burden in granulomas at mid and late timepoints. The reduction in bacterial burden also coincides with an increase in histologic pathologies relating to granuloma healing or resolution. Although the temporal presence of T-bet^+^ adaptive lymphocytes is likely driven by host and pathogen factors, vaccines and host-directed therapies aimed at facilitating faster recruitment of these functional adaptive T cells to *Mtb* infected lungs would likely promote bacterial control and containment of disease.

## ACKNOWLEDGEMENTS

We thank Cassaundra Ameel, Carolyn Bigbee, Ryan Kelly, Amy Fraser, Janelle Gleim, Abigail Gubernat, Kush Patel, Mark Rodgers, and Jennie Vorhauer for their laboratory and technical assistance; Beth Junecko for imaging assistance; Chelsea Causgrove, Dan Fillmore, Kara Kracinovsky, Skyler Pergalske, and Jenn Sakal for veterinary assistance; and other members of the Flynn, Scanga, and Lin laboratories for their collaborative efforts and insightful conversations. This study was supported by a T32 training grant NIH 5T32AI1065380-13 and NIH AI123093 (JLF and DEK).

## AUTHOR CONTRIBUTION

JLF and DK conceived of the study. NLG contributed to the acquisition, analysis, and interpretation of data for drafting of the manuscript. EK, PLL, LJF, and JT aided in implementing animal protocols, care, or specimen acquisition. EK provided histologic images and sample descriptions. HJB, AW, and PM contributed to PET CT analysis and PM provided statistical expertise. JTM provided immunofluorescence expertise. All authors revised and approved the final version of this manuscript.

## MATERIALS AND METHODS

### Ethics statement

All experiments, protocols, and care of animals were approved by the University of Pittsburgh School of Medicine Institutional Animal Care and Use Committee (IACUC). The Division of Laboratory Animal Resources and IACUC adheres to national guidelines established by the Animal Welfare Act (7 U.S. Code Sections 2131-2159) and the Guide for the Care and use of Laboratory Animals (Eighth Edition) as mandated by the U.S. Public Health Service Policy. Animals used in this study were housed in rooms with autonomously controlled temperature and provided enhanced enrichment procedures as previously described (Winchell et al., 2020).

### Animals and *Mtb* infection

Eight adult cynomolgus macaques (*Macaca fasicularis*) were infected with a low dose (15-20) CFU of *Mtb* (Erdman strain) via bronchoscopic installation as previously described (Supplemental Table 1) (Capuano et al., 2003b). NHPs were monitored daily in the Biosafety Level 3 (BSL3) laboratory at the University of Pittsburgh in compliance with the University’s Institutional Animal Care and Use Committee (IACUC). Prior to infection, animals were examined and placed in quarantine to evaluate health and confirm no prior *Mtb* infection. For pathology analysis and figures, banked control samples with corresponding PET CT scan data were used to supplement the animals dedicated to this study, with only original granulomas (i.e. those first observed on 4 week scans) being utilized (Supplemental Table 2).

### FDG PET CT imaging

Following *Mtb* infection, longitudinal PET CT imaging was performed to identify original granulomas and track disease over time. NHPs were sedated and injected with a PET tracer, 2-deoxy-2-(^18^F)Fluoro-D-glucose (FDG), and imaged using the Mediso MultiScan LFER 150 (Mediso, Budapest, Hungary) PET CT integrated preclinical scanner (White et al., 2017, Lin et al., 2013). Imaging was performed in accordance with biosafety and radiation safety requirements within the BSL3 facility at the University of Pittsburgh every 2-4 weeks beginning at 4 weeks post-infection until pre-determined animal endpoint. Scans were analyzed using OsiriX DICOM (Pixmeo, Geneva, Switzerland) viewer software by in-house trained PET CT analysts (Pauline Maiello, H. Jacob Borish, and Alexander G. White) (White et al., 2017, Rosset et al., 2004).

### Necropsy

Necropsy procedures were performed as previously described (Lin et al., 2009). In short, individual granulomas were identified using the pre-necropsy ^18^F-FDG PET CT scan and isolated along with lymph nodes and portions of uninvolved lung lobes. At necropsy, NHPs were sedated with ketamine, maximally bled and humanely euthanized using pentobarbital and phenytoin (Beuthanasia; Schering-Plough, Kenilworth, NJ). To assess gross pathology, animals were scored based on number, size, and pattern of lung granulomas and extent of disease involvement in lobes, mediastinal LNs, and visceral organs as previously described (Lin et al., 2009). Tissues were bisected and placed in formalin for paraffin embedding to perform histological evaluation. Single-cell suspensions were obtained for assessment of bacterial burden and immunological assays using gentle macs enzymatic dissociation (59% of samples) or physical homogenization. Bacterial burden was evaluated for individual tissue sections by plating serial dilutions of homogenate on 7H11 or PANTA agar plates and incubated at 37°C in 5%CO_2_ for 21 days. Total thoracic CFU is calculated from the summation of all lung, lung granuloma, and thoracic LN-plated samples (Maiello et al., 2017).

### Flow cytometry

Following processing, single cell suspensions underwent surface and TF/intracellular cytokine staining (ICS). Prior to staining, cells were incubated at 37°C in 5% CO_2_ in RPMI supplemented with 1%HEPES, 1% L-glutamine, 10% human AB serum, and 0.1% brefeldin A (Golgiplug; BD Biosciences, San Jose, CA) for 3 hours. Cells were stained with a viability dye (Zombie NIR) followed by surface stains (Supplemental Table 3) using standard protocols. TF and ICS was performed following permeabilization using True-Nuclear buffer kit according to the recommended protocol (True-Nuclear Transcription Factor Buffer Set; BioLegend, San Diego, CA). Samples were acquired on a Cytek Aurora (Cytek, Bethesda, MD) and analyzed using FlowJo Software (BD Biosciences) (Supplemental Fig. 2) and positive staining was verified against unstained controls. Only samples with >50 flow events in the parent population were reported. For analysis of CD3^-^CD20^-^ or CD20^+^ cells, animal 6319 was excluded due to poor staining with the anti-CD20 antibody. In total, 88 granuloma samples were taken for flow cytometric analysis (early=24, mid=31, and late=33) representing 94% (average) of the original lung granulomas isolated from animals at the time of necropsy (Supplemental Table 2).

### Histology

Individual tissue samples were formalin fixed paraffin embedded (FFPE) and cut into 5μm serial sections for tissue sectioning and histological evaluation. A veterinary pathologist (Edwin Klein) visually assessed the hematoxylin and eosin-stained lesions and described the histopathologic features and relevant cell types in each granuloma. Granuloma descriptions were categorized based on similar histopathologic description and analyzed to determine frequencies and presence of pathologic descriptors at each timepoint. Banked sections from animals with >3 granulomas having histologic descriptions were included for analyses.

### Immunofluorescence

Cut and mounted FFPE tissue sections were treated with xylenes twice, 5 minutes each, followed by graded ethanol (95%, 70%) incubations for deparaffinization. Slides were subsequently boiled with antigen retrieval buffer (Tris-EDTA, pH9, made in house or citrate buffer, pH6, Sigma C999-1000mL) and blocked for one hour with PBS containing 1% bovine serum albumin (BSA). Following blocking, slides were incubated with primary antibodies (Supplemental Table 3) for one hour at room temperature (RT) or 18 hours at 4°C in a humidified chamber. Fluorochrome-conjugated anti-mouse, rat, or rabbit antibodies, purchased from Jackson ImmunoResearch (Jackson ImmunoResearch Laboratories, West Grove, PA) and Thermo Fisher (Thermo Fisher, Waltham, MA), were used for secondary labeling for one hour at RT in a humidified chamber. Coverslips were mounted using ProLong Gold Antifade Mounting Medium with DAPI (Thermo Fisher) and imaged using an Olympus FV1000 confocal microscope (Olympus, Center Valley, PA) or Nikon e1000 (Nikon, Melville, NY) epifluorescent microscope. Post processing, images obtained on the Olympus confocal were stitched in Photoshop (Adobe Systems, Mountain View, CA) and images taken from both microscopes were brightened by applying a linear adjustment to the histogram levels for all channels in the entire image, taking care to maintain the integrity of the original image. Supplemental table 4 lists granulomas used for immunofluorescent analysis designated by figure.

### Statistical analysis and transformations

Adjusted cell counts were determined based on the hemocytometer count at the time of tissue homogenization (cells/mL) multiplied by the total volume of the sample. This value was then adjusted for sample splitting (for staining controls) and cutting (roughly half sent for histology). Lastly, a limit of detection value (LOD) determined by total volume of sample was added to all samples to account for those that fall below detectable levels. Kruskal-Wallis tests were performed to determine if there were differences in necropsy timepoint followed by Dunn’s multiple comparison tests to determine differences between specific groups. Non-parametric paired t tests were performed to compare frequencies of cytokine production from TF^+^ cells and TF^-^ cells within the same sample. For all tests, P values <0.05 were considered significant. CFU was log transformed (CFU+1/granuloma) to eliminate any zeroes from the analysis. Statistical analyses were performed using GraphPad Prism 9 (GraphPad Software, San Diego, CA).

**Supplementary Figure 1: Pathology and CFU in original granulomas**. (A) A range of granuloma pathologies is seen in original lesions across timepoints post-infection. (B) Individual granuloma CFU from fibrotic and non-fibrotic original granulomas isolated at mid and late timepoints. Statistics were performed using a Mann-Whitney test with p values noted on graph. (C) The proportion of sterile granulomas at each timepoint with total granuloma numbers assessed listed below (left). The proportion of fibrotic, neutrophilic, collagenic, or necrotic of the sterile granulomas from all timepoints (11 total)(right).

**Supplementary figure 2: Flow cytometry gating strategy for lung granulomas** (A) Identification of live events, singlets, and myeloid and lymphoid populations. (B) From the lymphocyte parent gate, identification of B cells, innate lymphocytes (ILCs) and T cells using CD3^+^ and CD20^+^. Examples of transcription factor, activation marker, and pro-inflammatory cytokine gating. (C) From the CD3^+^ parent gate, identification of CD4^+^, CD4^+^CD8^+^, CD8^+^, and CD4^-^CD8^-^ lymphocytes. Example of gating on transcription factor^+^ cells versus transcription factor^-^ cells and then gating on pro-inflammatory cytokines. (D) From the myeloid gate, an example of CD11b versus CD11c to determine single, negative, and co-expressing populations. Examples of single identification of CD11c, CD11b, and CD163 from the myeloid parent gate.

**Supplementary figure 3: Mid and late granulomas have lower proportions of CD11c**^**+**^ **and CD11b**^**+**^ **myeloid cells but higher levels of IL-10**. (A) Frequency and numbers of myeloid cells in granulomas from each timepoint. (B) Relative frequency of the combinations of CD11c and CD11b expression in myeloid cells from each timepoint where each bar represents a granuloma and animals are represented by horizontal bars (Fig. 1B). (C) Frequencies of CD11b, CD11c, and CD163 expression in myeloid cells across timepoints. (D) Cytokine expression in myeloid cells across timepoints. For A, C and D, individual points represent granulomas and animals are denoted by color. For A, C and D statistics, Kruskal Wallis tests were performed with Dunn’s multiple comparisons-adjusted p values reported on graphs.

**Supplementary figure 4: Lymphocytes produce low levels of pro-inflammatory cytokines at all timepoints despite changes in activation marker expression**. Pro-inflammatory cytokine and activation marker expression in (A) lymphocytes, (B) T cells: CD3^+^CD4^+^, CD3^+^CD4^+^CD8+, CD3^+^CD8^+^, and (C) CD3^-^CD20^-^ cells. Kruskal Wallis tests were performed with Dunn’s multiple comparisons-adjusted p values reported on graphs.

**Supplementary figure 5: Transcription factor**^**+**^ **lymphocyte subpopulations in original granulomas over time**. Single or double transcription factor expression in (A) CD4^+^, (B) CD3^+^CD4^+^CD8^+^, (C) CD3^+^CD8^+^, (D) CD3^+^CD4^-^CD8^-^, (E) CD20^+^, (F) CD3^-^CD20^-^ cells. (G) Frequency of dual expression of T-bet and RORa (left) or T-bet and GATA3 (right) in all lymphocytes. Kruskal Wallis tests were performed with Dunn’s multiple comparisons-adjusted p values reported on graphs.

**Supplementary figure 6: Immunofluorescence staining for RORα, GATA3, Foxp3, and Granzyme B in granulomas**. (A) Immunofluorescence for RORα (red, arrows in inset) in an early timepoint granuloma can be observed (although rare) within CD3^+^ (green) cells. (B) Immunofluorescence for GATA3 (red, arrows in inset) in a late timepoint granuloma can be observed (although rare) within CD3^+^ (green) cells. (C) Foxp3 (blue) within CD3^+^ (green) CD4^+^ (red) cells in a mid timepoint granuloma. (D) Granzyme B immunofluorescence staining in a late timepoint granuloma is observed within CD3^+^ (green) Tbet^+^ (blue) cells as indicated by arrows.

**Supplementary figure 7: Pro-inflammatory cytokine expression is higher in T-bet**^**+**^ **CD3**^**-**^ **CD20**^**-**^ **cells**. Frequency of IFN-γ and TNF in T-bet^+^ or T-bet^-^ CD3^-^CD20^-^ cells in granulomas from the 4 week timepoint. Statistics performed using Wilcoxon signed-rank with p values reported on graph.

**Supplementary table 1: Animal and *Mtb* infection data**

**Supplementary table 2: Granulomas used for flow cytometry analysis**

**Supplementary table 3: List of antibodies used to identify cellular populations**

**Supplementary table 4: Granulomas used for IHC and immunofluorescence, designated by figure**

---

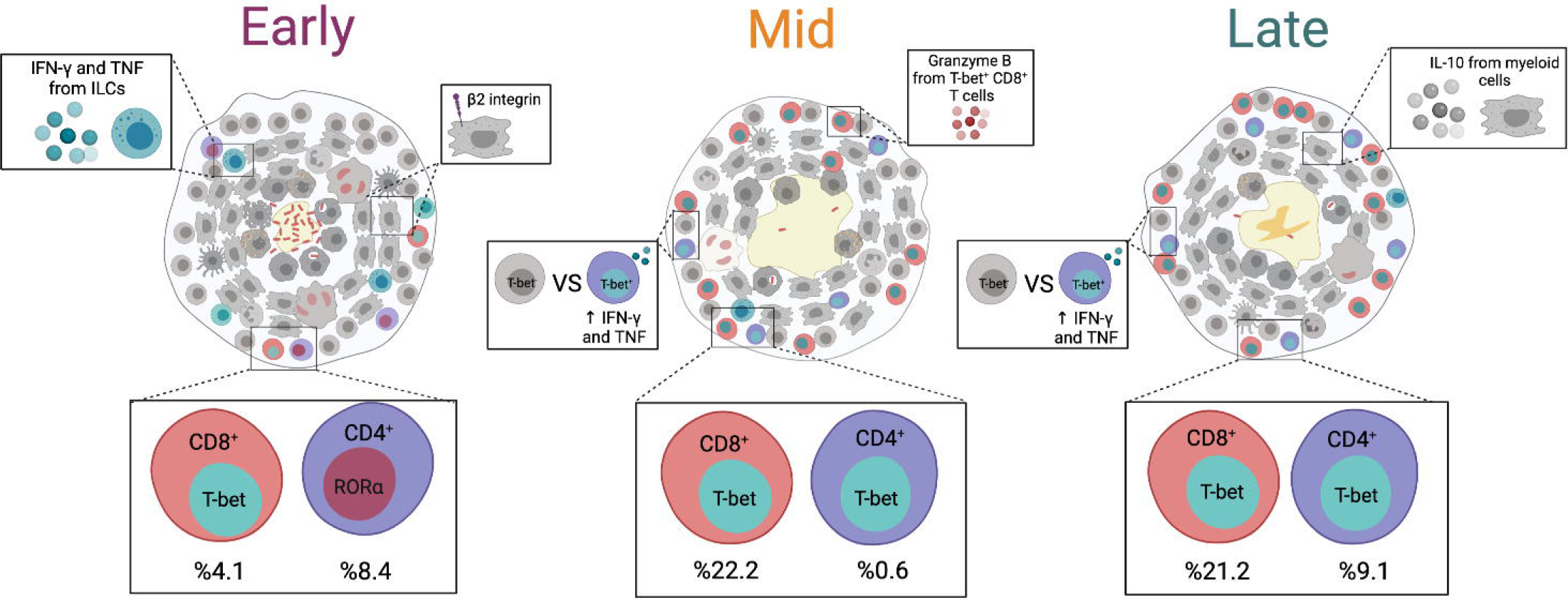

