## Supplemental Figures for "T cell transcription factor expression evolves as adaptive immunity matures in granulomas from *Mycobacterium tuberculosis*-infected cynomolgus macaques"

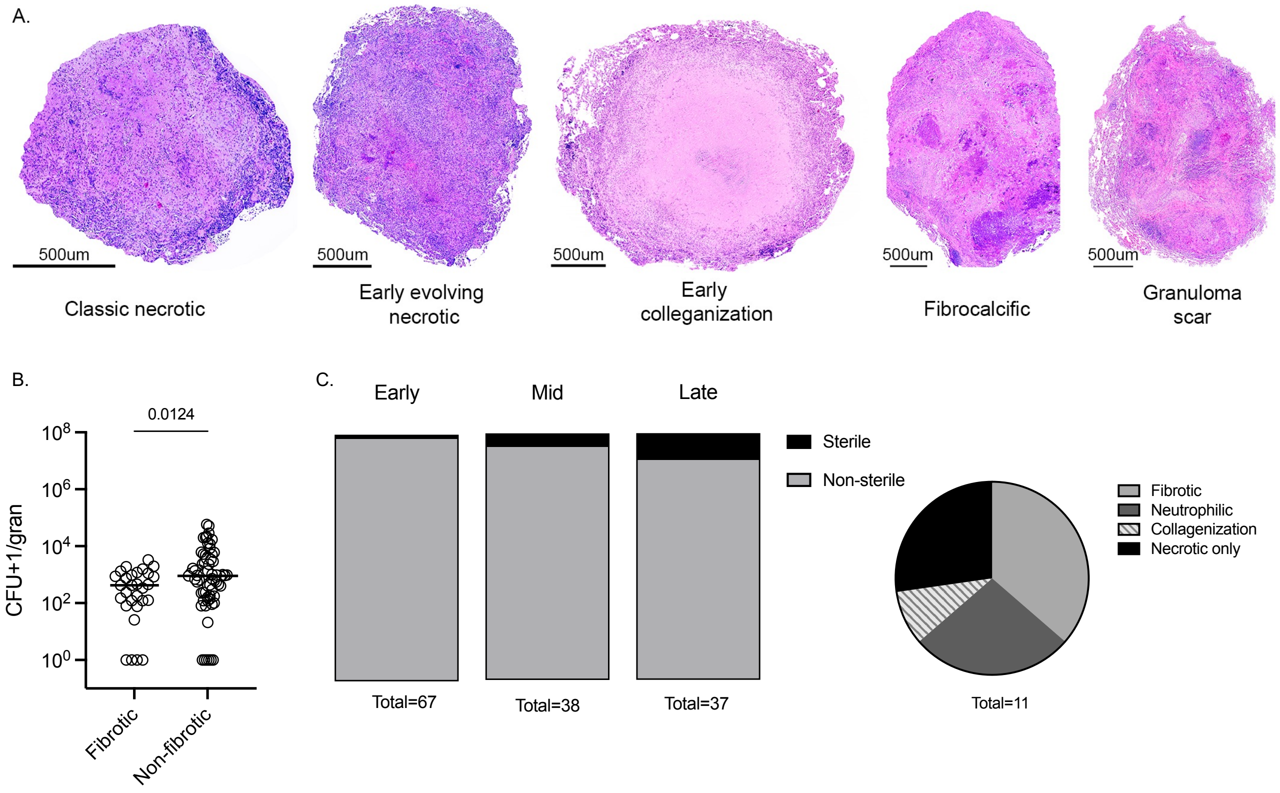
A.
500um
Classic necrotic
Early evolving necrotic
Early colleganization
Fibrocalcific
Granuloma scar
B.
0.0124
CFU+1/gran
$10^8$
$10^6$
$10^4$
$10^2$
$10^0$
Fibrotic
Non-fibrotic
C.
Early
Mid
Late
Sterile
Non-sterile
Total=67
Total=38
Total=37
Fibrotic
Neutrophilic
Collagenization
Necrotic only
Total=11

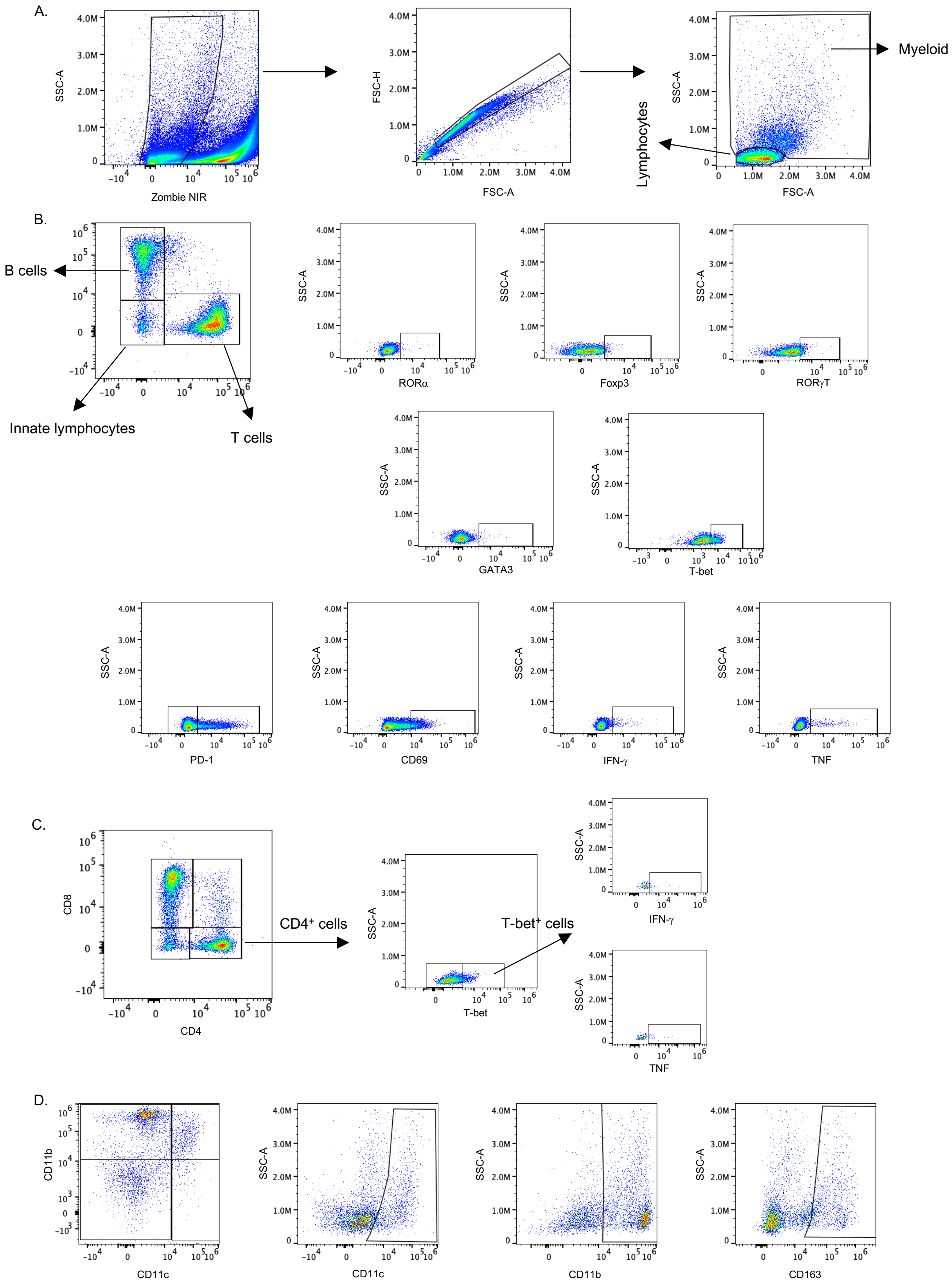

A.
SSC-A
Zombie NIR
FSC-H
FSC-A
Lymphocytes
Myeloid
B.
B cells
Innate lymphocytes
T cells
RORα
Foxp3
RORγT
GATA3
T-bet
PD-1
CD69
IFN-γ
TNF
C.
CD8
CD4
CD4⁺ cells
T-bet⁺ cells
D.
CD11b
CD11c
CD163

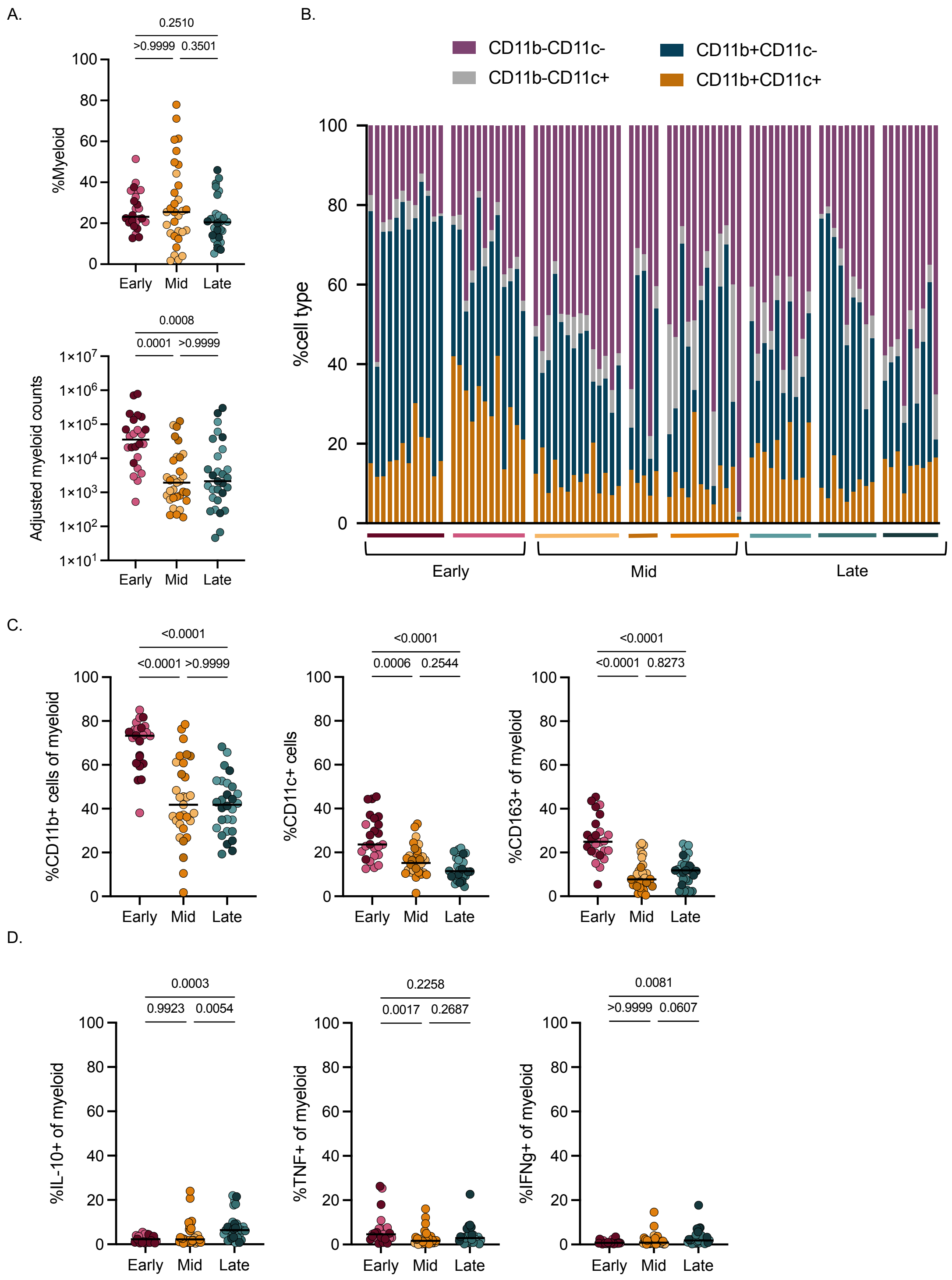

A.
0.2510
>0.9999
0.3501
%Myeloid
Early
Mid
Late
0.0008
0.0001
>0.9999
Adjusted myeloid counts
B.
CD11b-CD11c-
CD11b-CD11c+
CD11b+CD11c-
CD11b+CD11c+
%cell type
Early
Mid
Late
C.
<0.0001
<0.0001
>0.9999
%CD11b+ cells of myeloid
0.0006
0.2544
%CD11c+ cells
<0.0001
0.8273
%CD163+ of myeloid
D.
0.0003
0.9923
0.0054
%IL-10+ of myeloid
0.2258
0.0017
0.2687
%TNF+ of myeloid
0.0081
>0.9999
0.0607
%IFNg+ of myeloid

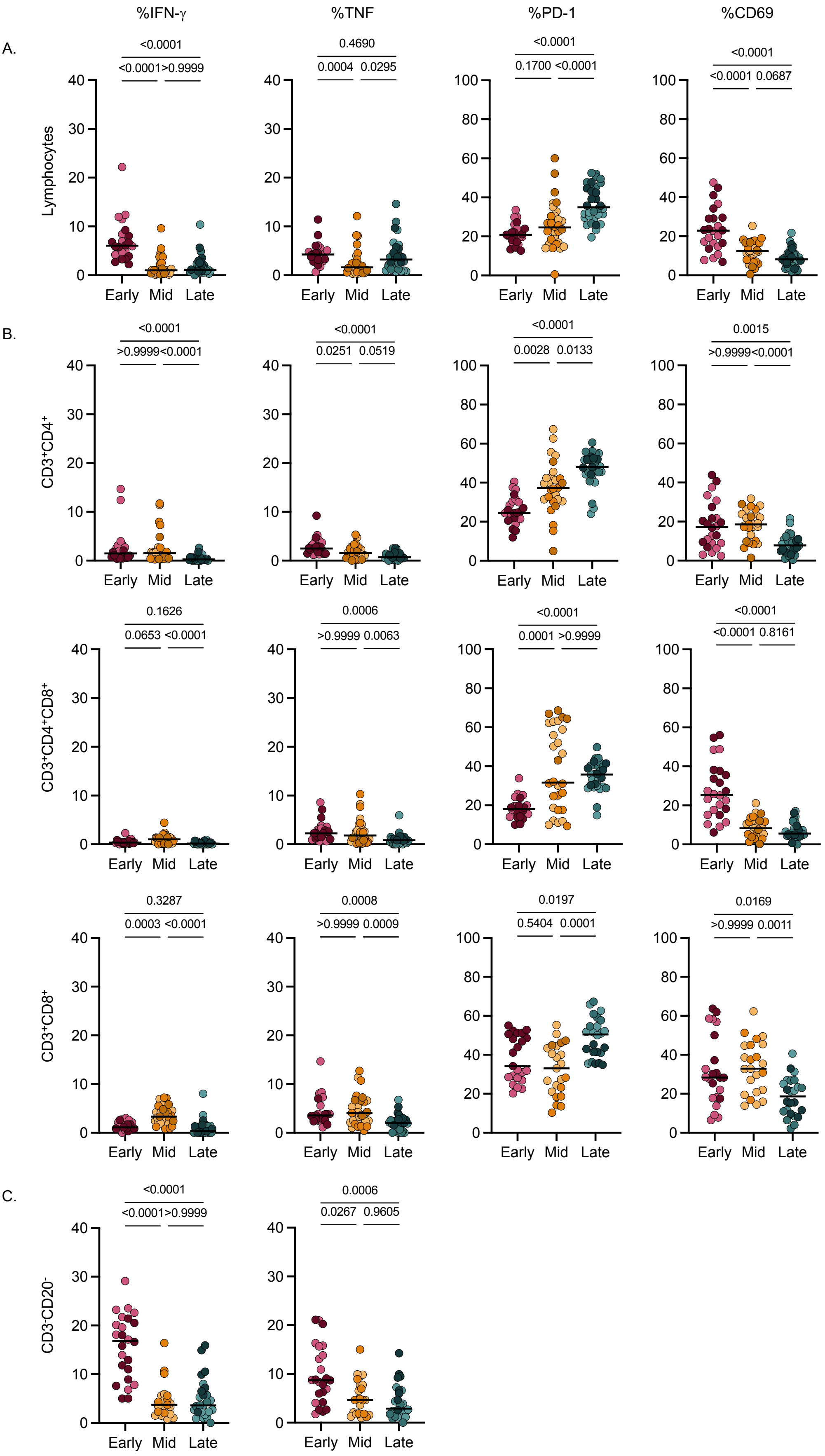
%IFN-γ
%TNF
%PD-1
%CD69
A.
B.
C.
Lymphocytes
CD3+CD4+
CD3+CD4+CD8+
CD3+CD8+
CD3-CD20-
Early
Mid
Late
<0.0001
<0.0001
>0.9999
0.4690
0.0004
0.0295
<0.0001
0.1700
<0.0001
<0.0001
<0.0001
0.0687
<0.0001
>0.9999
<0.0001
<0.0001
0.0251
0.0519
<0.0001
0.0028
0.0133
0.0015
>0.9999
<0.0001
0.1626
0.0653
<0.0001
0.0006
>0.9999
0.0063
<0.0001
0.0001
>0.9999
<0.0001
<0.0001
0.8161
0.3287
0.0003
<0.0001
0.0008
>0.9999
0.0009
0.0197
0.5404
0.0001
0.0169
>0.9999
0.0011
<0.0001
<0.0001
>0.9999
0.0006
0.0267
0.9605

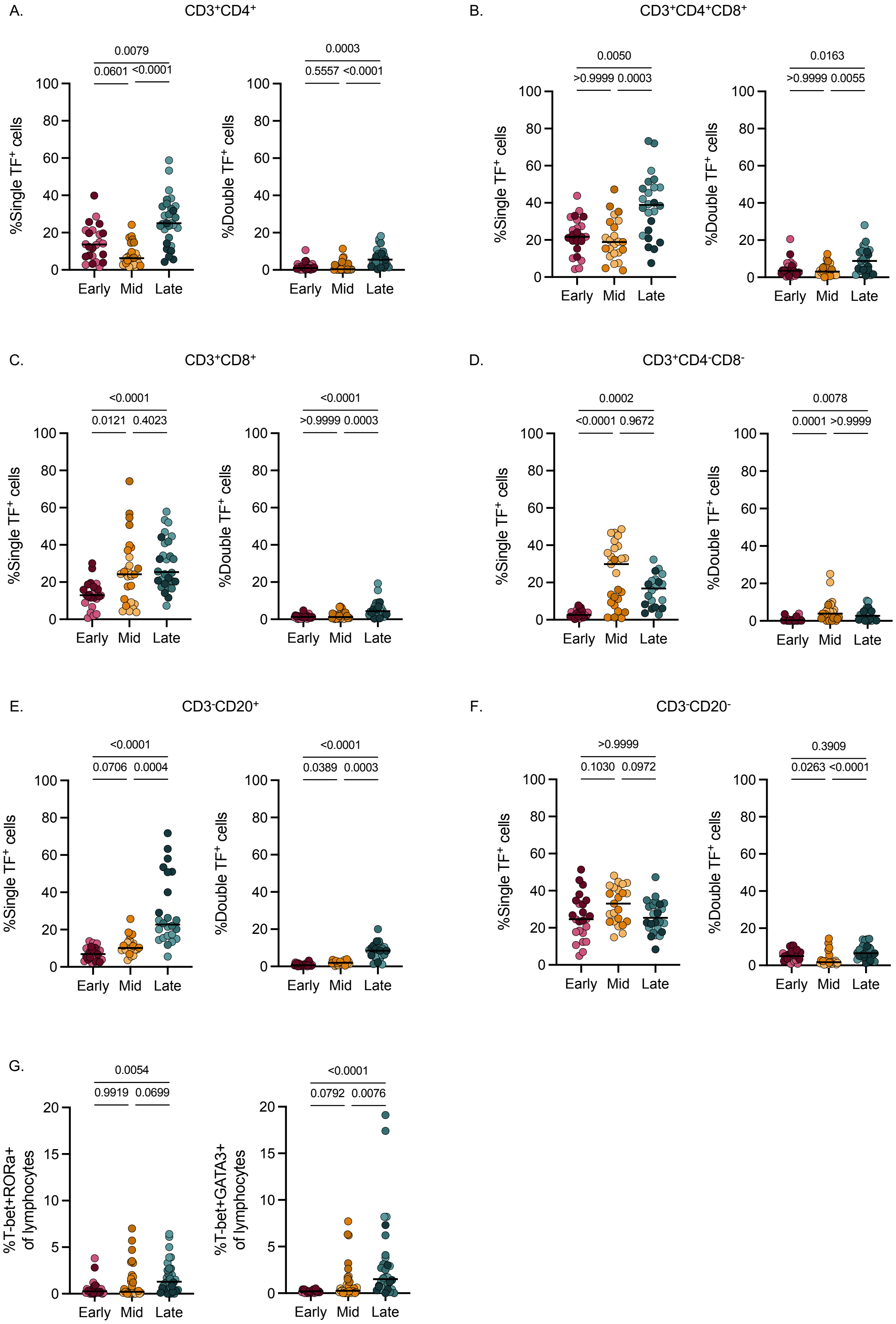
A.
CD3+CD4+
0.0079
0.0601 <0.0001
%Single TF+ cells
Early Mid Late
0.0003
0.5557 <0.0001
%Double TF+ cells
Early Mid Late
B.
CD3+CD4+CD8+
0.0050
>0.9999 0.0003
%Single TF+ cells
Early Mid Late
0.0163
>0.9999 0.0055
%Double TF+ cells
Early Mid Late
C.
CD3+CD8+
<0.0001
0.0121 0.4023
%Single TF+ cells
Early Mid Late
<0.0001
>0.9999 0.0003
%Double TF+ cells
Early Mid Late
D.
CD3+CD4-CD8-
0.0002
<0.0001 0.9672
%Single TF+ cells
Early Mid Late
0.0078
0.0001 >0.9999
%Double TF+ cells
Early Mid Late
E.
CD3-CD20+
<0.0001
0.0706 0.0004
%Single TF+ cells
Early Mid Late
<0.0001
0.0389 0.0003
%Double TF+ cells
Early Mid Late
F.
CD3-CD20-
>0.9999
0.1030 0.0972
%Single TF+ cells
Early Mid Late
0.3909
0.0263 <0.0001
%Double TF+ cells
Early Mid Late
G.
0.0054
0.9919 0.0699
%T-bet+RORa+ of lymphocytes
Early Mid Late
<0.0001
0.0792 0.0076
%T-bet+GATA3+ of lymphocytes
Early Mid Late

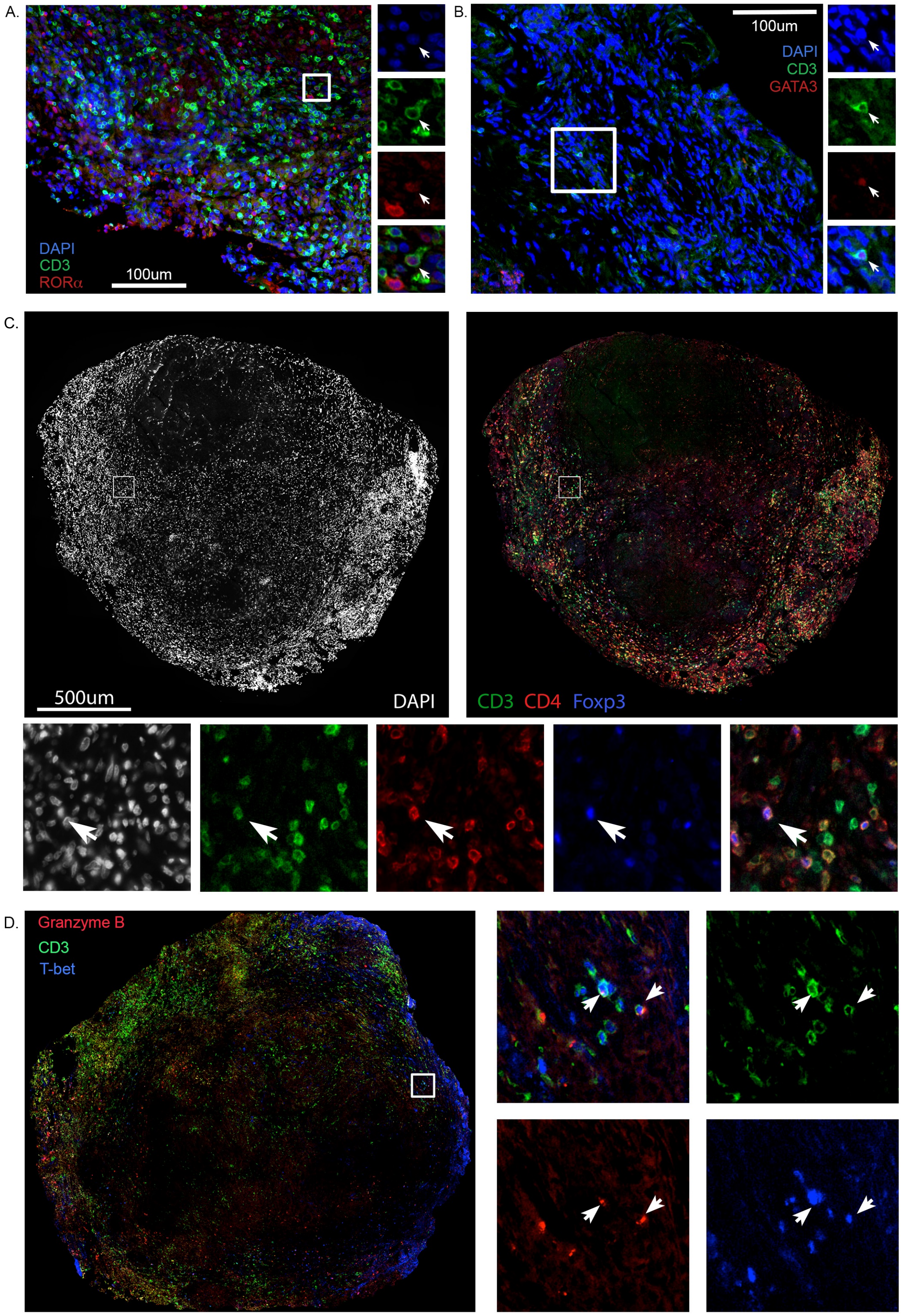
A.
DAPI
CD3
RORα
100um
B.
100um
DAPI
CD3
GATA3
C.
500um
DAPI
CD3 CD4 Foxp3
D.
Granzyme B
CD3
T-bet

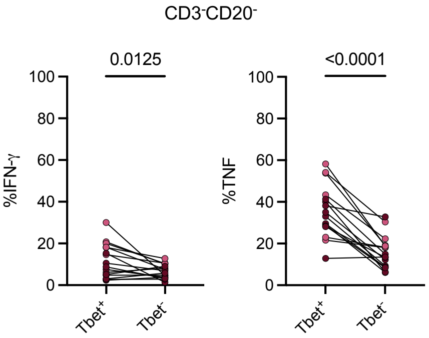
CD3-CD20-
0.0125
<0.0001
%IFN-γ
%TNF
100
80
60
40
20
0
Tbet+
Tbet-

| Animal | Necropsy group | Necropsy timepoint | CFU @ infection | Date of infection | Date of Necropsy | # Lung granulomas | # Original granulomas | # Original granulomas for flow | Total CFU | Lung CFU | LN CFU | Necropsy score | Total Pet Hot |
|---|---|---|---|---|---|---|---|---|---|---|---|---|---|
| 32719 | Early | 4 weeks | 19 | 12/17/19 | 1/13/20 | 12 | 12 | 12 | 5.61E+06 | 4.40E+06 | 1.21E+06 | 21 | 22449.25 |
| 32819 | Early | 4 weeks | 19 | 12/17/19 | 1/13/20 | 12 | 12 | 12 | 7.34E+05 | 5.96E+05 | 1.38E+05 | 20 | 8789.65 |
| 6219 | Mid | 12 weeks | 10 | 7/17/19 | 10/9/19 | 14 | 17 | 14 | 1.51E+05 | 1.11E+05 | 4.05E+04 | 27 | 16627.7 |
| 6319 | Mid | 12 weeks | 10 | 7/17/19 | 10/7/19 | 14 | 5 | 5 | 3.73E+04 | 2.44E+04 | 1.29E+04 | 25 | 4467.49 |
| 6419 | Mid | 12 weeks | 10 | 7/17/19 | 10/9/19 | 19 | 12 | 12 | 2.98E+04 | 6.37E+03 | 2.34E+04 | 23 | 1253.85 |
| 6519 | Late | 20 weeks | 18 | 8/22/19 | 1/6/20 | 19 | 14 | 12 | 1.23E+04 | 1.18E+04 | 5.50E+02 | 16 | 620.61 |
| 6619 | Late | 20 weeks | 18 | 8/22/19 | 1/6/20 | 12 | 11 | 11 | 1.57E+04 | 4.15E+03 | 1.15E+04 | 23 | 2663.83 |
| 6819 | Late | 20 weeks | 18 | 8/22/19 | 1/8/20 | 14 | 11 | 10 | 2.90E+04 | 1.87E+04 | 1.03E+04 | 24 | 6167.06 |

| Animal | Sample | NX Timepoint (wks) | NX Time | Gentle macs processed | CFU/ granuloma |
|---|---|---|---|---|---|
| 32819 | RLL 1 (GM) | 4 | Early | Yes | 18600 |
| 32819 | RLL 2 | 4 | Early | No | 520 |
| 32819 | RLL 3 (GM) | 4 | Early | Yes | 46200 |
| 32819 | RLL 7 (GM) | 4 | Early | Yes | 3120 |
| 32819 | RLL 4 | 4 | Early | No | 30000 |
| 32819 | RL 5 (GM) | 4 | Early | Yes | 5580 |
| 32819 | RLL 6 (GM) | 4 | Early | Yes | 28800 |
| 32819 | RLL 8 | 4 | Early | No | 103950 |
| 32819 | RLL 9 (GM) | 4 | Early | Yes | 276000 |
| 32819 | RLL 10 (GM) | 4 | Early | Yes | 71400 |
| 32819 | RLL 11 | 4 | Early | No | 3360 |
| 32819 | RLL 13 | 4 | Early | No | 1880 |
| 32719 | LLL gr 12 | 4 | Early | No | 930000 |
| 32719 | LLL gr 2 | 4 | Early | No | 40950 |
| 32719 | LLL gr 3/4 | 4 | Early | No | 45500 |
| 32719 | LLL gr 6 (GM) | 4 | Early | Yes | 762000 |
| 32719 | LLL gr 7 (GM) | 4 | Early | Yes | 1512000 |
| 32719 | LLL gr 9 | 4 | Early | No | 91200 |
| 32719 | LLL gr 10 (GM) | 4 | Early | Yes | 96000 |
| 32719 | LLL gr 13 | 4 | Early | No | 76800 |
| 32719 | LLL gr 14 (GM) | 4 | Early | Yes | 480000 |
| 32719 | LLL gr 15 (GM) | 4 | Early | Yes | 70200 |
| 32719 | LLL gr 16 (GM) | 4 | Early | Yes | 135600 |
| 32719 | LLL gr 17 (GM) | 4 | Early | Yes | 93000 |
| 6219 | RLL cluster 6 | 12 | Mid | No | 1900 |
| 6219 | RLL cluster 13 | 12 | Mid | No | 3750 |
| 6219 | RLL gran 1 (GM) | 12 | Mid | Yes | 635 |
| 6219 | RLL gran 3 (GM) | 12 | Mid | Yes | 1360 |
| 6219 | RLL gran 5 (GM) | 12 | Mid | Yes | 875 |
| 6219 | RLL gran 9 (GM) | 12 | Mid | Yes | 450 |
| 6219 | RLL gran 10 (GM) | 12 | Mid | Yes | 605 |
| 6219 | RLL gran 11 (GM) | 12 | Mid | Yes | 990 |
| 6219 | RLL gran 12 (GM) | 12 | Mid | Yes | 650 |
| 6219 | RLL gran 14 | 12 | Mid | No | 975 |
| 6219 | RLL gran 16 | 12 | Mid | No | 900 |
| 6219 | RLL gran 17 | 12 | Mid | No | 960 |
| 6219 | RLL gran 19 | 12 | Mid | No | 400 |
| 6219 | RLL gran 22 (GM) | 12 | Mid | Yes | 1355 |
| 6319 | RLL cluster 5/6 | 12 | Mid | No | 975 |
| 6319 | RLL gran 7 (GM) | 12 | Mid | No | 7080 |
| 6319 | RLL gran 9 (GM) | 12 | Mid | Yes | 6000 |
| 6319 | RLL gran 18 (GM) | 12 | Mid | Yes | 0 |
| 6319 | RLL gran 19 (GM) | 12 | Mid | Yes | 6150 |
| 6419 | LLL gran 1 (GM) | 12 | Mid | Yes | 25 |
| 6419 | LLL gran 2 | 12 | Mid | No | 1325 |
| 6419 | LLL gran 3 | 12 | Mid | No | 700 |
| 6419 | LLL gran 4 (GM) | 12 | Mid | Yes | 335 |
| 6419 | LLL gran 5 (GM) | 12 | Mid | Yes | 120 |
| 6419 | LLL gran 6A | 12 | Mid | No | 1980 |
| 6419 | LLL gran 7 (GM) | 12 | Mid | Yes | 440 |
| 6419 | LLL gran 8 | 12 | Mid | No | 50 |
| 6419 | LLL gran 9 (GM) | 12 | Mid | Yes | 115 |
| 6419 | LLL gran 11 (GM) | 12 | Mid | Yes | 470 |
| 6419 | LLL gran 12 | 12 | Mid | No | 120 |
| 6419 | LLL gran E | 12 | Mid | No | 80 |
| 6519 | RLL cluster 12 | 20 | Late | No | 560 |
| 6519 | RLL gran H | 20 | Late | No | 50 |
| 6519 | RLL gran 1 | 20 | Late | No | 10 |
| 6519 | RLL gran 3 (GM) | 20 | Late | Yes | 870 |
| 6519 | RLL gran 4 (GM) | 20 | Late | Yes | 6030 |
| 6519 | RLL gran 5 (GM) | 20 | Late | Yes | 1105 |
| 6519 | RLL gran 9 (GM) | 20 | Late | Yes | 750 |
| 6519 | RLL gran 13 (GM) | 20 | Late | Yes | 975 |
| 6519 | RLL gran 14 | 20 | Late | No | 0 |
| 6519 | RLL gran 16 (GM) | 20 | Late | Yes | 180 |
| 6519 | RLL gran 17 (GM) | 20 | Late | Yes | 300 |
| 6519 | RLL gran 20 | 20 | Late | No | 0 |
| 6619 | LLL cluster 2 | 20 | Late | No | 450 |
| 6619 | LLL gran 1 (GM) | 20 | Late | Yes | 240 |
| 6619 | LLL gran 3 | 20 | Late | No | 850 |
| 6619 | LLL gran 4 (GM) | 20 | Late | Yes | 320 |
| 6619 | LLL gran 5 | 20 | Late | No | 10 |
| 6619 | LLL gran 6 | 20 | Late | No | 75 |
| 6619 | LLL gran 7 old (GM) | 20 | Late | Yes | 175 |
| 6619 | LLL gran 8 (GM) | 20 | Late | Yes | 240 |
| 6619 | LLL gran 9 old (GM) | 20 | Late | Yes | 1183 |
| 6619 | LLL gran 10 (GM) | 20 | Late | Yes | 180 |
| 6619 | LLL gran 11 old (GM) | 20 | Late | Yes | 425 |
| 6819 | LLL gran 1 | 20 | Late | No | 400 |
| 6819 | LLL gran 5 (GM) | 20 | Late | Yes | 5285 |
| 6819 | LLL gran 7 (GM) | 20 | Late | Yes | 2280 |
| 6819 | LLL gran 8 (GM) | 20 | Late | Yes | 930 |
| 6819 | LLL gran 9 (GM) | 20 | Late | Yes | 1015 |
| 6819 | LLL gran 10 | 20 | Late | No | 100 |
| 6819 | LLL gran 12 (GM) | 20 | Late | Yes | 720 |
| 6819 | LLL gran 14 (GM) | 20 | Late | Yes | 1650 |
| 6819 | LLL gran 15 (GM) | 20 | Late | Yes | 2940 |
| 6819 | LLL gran 18 | 20 | Late | No | 260 |

| Target | Clone | Company | Catalog # | Method |
|---|---|---|---|---|
| CD69 | TP1.55.3 | Beckman Coulter | 6607110 | Flow |
| ROR$\alpha$ | NR1F1 | R&D | IC8924P-100 | Flow |
| CD163 | GHI/61 | BioLegend | 333614 | Flow |
| Foxp3 | PCH101 | Invitrogen | 45-4776-42 | Flow |
| ROR$\gamma$T | AFKJS-9 | Thermo fisher | 46-6988-80 | Flow |
| T-bet | 4B10 | BioLegend | 644812 | Flow |
| IFN-g | B27 | BioLegend | 562974 | Flow |
| CD4 | L200 | BD | 563094 | Flow |
| CD11c | 3.9 | BioLegend | 301628 | Flow |
| PD-1 (CD279) | EH12.1 | BD | 566175 | Flow |
| IL-10 | JES3-9D7 | eBioscience | 48-7108-42 | Flow |
| TNF | Mab11 | BioLegend | 563418 | Flow |
| CD3 | SP34-2 | BD | 624072 | Flow |
| GATA3 | 16E10A23 | BioLegend | 653806 | Flow |
| CD20 | 2H7 | BioLegend | 302318 | Flow |
| CD8 | RPA-T8 | BD | 563795 | Flow |
| CD11b | ICRF44 | BD | 741357 | Flow |
| ROR$\alpha$ | polyclonal | abcam | ab60134 | IHC |
| GATA-3 | EPR16651 | abcam | ab199428 | IHC |
| T-bet | D6N8B | Cell Signaling Technology | 13232 | IHC |
| Foxp3 | 236A/E7 | ebioscience | 14-4777-82 | IHC |
| CD163 | 10D6 | Neomarkers (thermo fisher) | MS-1103-S1 | IHC |
| CD11c | 5D11 | Novocastra- Leica | NCL-L-CD11c-563 | IHC |
| CD3 | CD3-12 | abcam | ab11089 | IHC |
| Granzyme B | GrB-7 | Dako | M7235 | IHC |
| CD8$\alpha$ | D8A8Y | CST | 85336S | IHC |
| CD3 | polyclonal | Dako | A0452 | IHC |

| Animal ID | Sample name | NX timing | CFU | Figure 1 | Figure 5 | Supplementary Figure 2 | Supplementary Figure 6 |
|---|---|---|---|---|---|---|---|
| 17313 | LLL gran 25A | Early | 1.2E+04 | D | | | |
| 9814 | RLL gran 12 | Mid | 4.6E+02 | D | A | | Foxp3 |
| 20512 | RLL gran 1 | Late | 0.0E+00 | D | | | |
| 16213 | RLL gran 16 | Late | 0.0E+00 | | | | GATA3 |
| 11114 | RLL gran 5 | Early | 5.0E+04 | | | | RORα |
| 15312 | Access gran 1 | Late | 2.4E+02 | | | | Granzyme B |
| 6619 | LLL gran 9 | Late | 1.2E+03 | | | Fibrocalcific | |
| 6619 | LLL gran 7 | Late | 1.8E+02 | | | Granuloma scar | |
| 13118 | RLL gran 7 | Early | 3.5E+04 | | | Early evolving necrotic | |
| 6219 | RLL gran 5 | Mid | 8.8E+02 | | | Classic necrotic | |
| 9814 | RLL gran 28 | Mid | 9.0E+02 | | | Early colleganization | |
| 9714 | RLL gran 27 | Mid | 5.4E+02 | | B | | |
